# Single-cell transcriptional atlas of the Chinese horseshoe bat (*Rhinolophus sinicus*) provides insight into the cellular mechanisms which enable bats to be viral reservoirs

**DOI:** 10.1101/2020.06.30.175778

**Authors:** Lili Ren, Chao Wu, Li Guo, Jiacheng Yao, Conghui Wang, Yan Xiao, Angela Oliveira Pisco, Zhiqiang Wu, Xiaobo Lei, Yiwei Liu, Leisheng Shi, Lianlian Han, Hu Zhang, Xia Xiao, Jingchuan Zhong, Hongping Wu, Mingkun Li, Stephen R. Quake, Yanyi Huang, Jianbin Wang, Jianwei Wang

**Affiliations:** NHC Key Laboratory of Systems Biology of Pathogens and Christophe Mérieux Laboratory, Institute of Pathogen Biology, Chinese Academy of Medical Sciences & Peking Union Medical College, Beijing 100730, China; Key Laboratory of Respiratory Disease Pathogenomics, Chinese Academy of Medical Sciences and Peking Union Medical College, Beijing, 100730, China; School of Life Sciences, Tsinghua-Peking Center for Life Sciences, Tsinghua University, Beijing 100084, China; Chan Zuckerberg Biohub, San Francisco, CA 94158, USA; NHC Key Laboratory of Systems Biology of Pathogens, Institute of Pathogen Biology, Chinese Academy of Medical Sciences & Peking Union Medical College, Beijing 100730, China; Beijing Institute of Genomics, Chinese Academy of Sciences, and China National Center for Bioinformation, Beijing, 100101, China; Center for Excellence in Animal Evolution and Genetics, Chinese Academy of Sciences, Kunming, 650223, China; Departments of Bioengineering and Applied Physics, Stanford University, Stanford, CA 94305, USA; Biomedical Pioneering Innovation Center (BIOPIC), Beijing Advanced Innovation Center for Genomics (ICG), College of Chemistry, and Peking-Tsinghua Center for Life Sciences, Peking University, Beijing 100871, China; Institute for Cell Analysis, Shenzhen Bay Laboratory, Guangdong 518132, China; Beijing Advanced Innovation Center for Structural Biology (ICSB), Tsinghua University, Beijing 100084, China; Chinese Institute for Brain Research (CIBR), Beijing 102206, China

**Keywords:** bat, single-cell sequencing, immunity, receptor, zoonotic

## Abstract

Bats are a major “viral reservoir” in nature and there is a great interest in not only the cell biology of their innate and adaptive immune systems, but also in the expression patterns of receptors used for cellular entry by viruses with potential cross-species transmission. To address this and other questions, we created a single-cell transcriptomic atlas of the Chinese horseshoe bat (*Rhinolophus sinicus*) which comprises 82,924 cells from 19 organs and tissues. This atlas provides a molecular characterization of numerous cell types from a variety of anatomical sites, and we used it to identify clusters of transcription features that define cell types across all of the surveyed organs. Analysis of viral entry receptor genes for known zoonotic viruses showed cell distribution patterns similar to that of humans, with higher expression levels in bat intestine epithelial cells. In terms of the immune system, CD8+ T cells are in high proportion with tissue-resident memory T cells, and long-lived effector memory nature killer (NK) T-like cells (*KLRG1*, *GZMA* and *ITGA4* genes) are broadly distributed across the organs. Isolated lung primary bat pulmonary fibroblast (BPF) cells were used to evaluate innate immunity, and they showed a weak response to interferon β and tumor necrosis factor-α compared to their human counterparts, consistent with our transcriptional analysis. This compendium of transcriptome data provides a molecular foundation for understanding the cell identities, functions and cellular receptor characteristics for viral reservoirs and zoonotic transmission.

## Introduction

Bats function as natural viral reservoirs and are distributed globally; they are unique as flying mammals. They have high diversity with more than 1,300 bat species having been identified [http://www.batcon.org]^1,2^. Bats carry some of the deadliest viruses for humans, including lyssaviruses, Ebola (EBOV) and Marburg (MARV) filoviruses, severe acute respiratory syndrome coronaviruses (SARS-CoV)-like viruses (SL-CoVs), Middle East respiratory syndrome (MERS-CoV)-like viruses (ML-CoVs), Hendra (HeV) and Nipah (NiV) henipaviruses^3,4^. SARS-CoV-2, which emerged in December 2019 and caused a global pandemic, is also considered as originating in bats^5,6^.

Bats have evolved over eons to sustain infection from pathogens without succumbing to overt disease, which indicates a uniquely powerful immune system^7^. According to the comparative genome and transcriptome studies, *in vitro* bat cell culture, and experimental infection assays, the diverse bat species may have evolved different mechanisms to balance between enhanced immune function which clears viral infections and tolerance on limiting immunopathology^1,8^. However, knowledge of bat immunology is still poorly understood as current studies used mainly *Pteropus alecto*, *Myotis davidii*, and *Rousettus aegyptiacus* species, but obtained conflicting findings on the function of bat immune systems^9,10^. The natural killer (NK) cells and type I interferons (IFNs) signaling pathways are of great interest. It has been reported a few NK cell receptor genes, killer cell lectin-like receptor genes (*KLRD* and *KLRC*), exist in the *Pteropus alecto* transcriptome and *Rousettus aegyptiacus* genome^11^. However, the majority of known canonical NK cell receptor genes are absent in currently known bat genomes^10,12^.

As they have a long life span and continued natural selection, the bat has also been considered as an excellent model to study human cellular evolution features compared to other lab animals^1^. Characterizing the extent to which bat cellular biological functions mirrors those of humans will enable scientists to understand the characteristics of the immune system and mechanisms of the zoonotic virus spreading. The exploration of the organs in single cell level in human, model mouse has provided insights into cellular diversity and revealed new cell types related to physiological function^13–15^.

In this study we report the molecular composition of 89 cell types from the Chinese horseshoe bat (*Rhinolophus sinicus*), belonging to suborder of *Yinpterochiroptera*, a natural reservoir of SL-CoVs. The compendium comprises single-cell transcriptomic data from cells of 19 organs, including adipose tissues (brown and white), bladder, bone marrow, brain, heart, intestine, kidney, liver, lung, muscle, pancreas, wing membrane, spleen, testis, thymus, tongue, trachea and whole blood.

## Results

### Transcriptomic characteristics

To ensure accuracy of the single cell sequencing (sc-seq), 12 of the 19 obtained organs were firstly analyzed by using bulk sequencing (bulk-seq), including adipose tissues (brown and white), brain, heart, intestine, kidney, liver, lung, muscle, spleen, tongue, and trachea (Fig. 1a). As the interaction of the virus with its cellular receptor is a key step in its pathogenesis^16^, we first compared the transcriptomic patterns of viral receptor genes in different mammals: bat, human and mouse. We analyzed 6 out of 28 known human viral receptor genes as representatives (Fig. 1b), in which five are known bat zoonotic virus receptors: angiotensin convert enzyme 2 (ACE2) (receptor of SARS-CoV, SARS-CoV-2 and human coronavirus (HCoV) NL63), dipeptidyl peptidase 4 (DPP4), (receptor of MERS-CoV), aminopeptidase N (ANPEP) (receptor of HCoV-229E)^17^, Ephrin-B2 (EFNB2) (receptor of HeV and NiV), NPC intracellular cholesterol transporter 1 (NPC1) (receptor of EBOV and MARV), coxsackievirus and adenovirus receptor [CXADR, the human and bat adenovirus (Adv) shared receptor]^18^. Transmembrane serine protease 2 (TMPRSS2), a protease essential for SARS-CoV and SARS-CoV-2 entry was also analyzed (Fig. 1b). The data show that *TMPRSS2, DPP4* and *ANPEP* express at high levels in the lung, intestine and kidney in bat, human and mouse. Although *ACE2* is also highly expressed in the intestine and kidney of all three species, for lung it expresses highly in only mouse but not in human or bat (Fig. 1b, 1c). Both the *ACE2* and *TMPRSS2* genes highly express in bat tongue, while only *ACE2* highly expresses in mouse tongue and only *TMPRSS2* in human tongue. In the heart, brain, spleen, wing (skin), muscle and adipose tissues, only *ACE2* shows highly expressed in two of the species, but *TMPRSS2* shows low expression levels. The neural cell adhesion molecule 1 (NCAM1) gene, Rabies virus (RABV) receptor, is found mainly expressed in the brain of all three species, as well as in the human heart, spleen and muscle. The EFNB2 and NPC1 are distributed in most of the organs in bat, human and mouse with similar expression level. Such distribution characteristics may be related to the multiple organs involved infections of HeV, NiV and EBOV^19,20^. Of the other 20 viral receptor genes, we notice that most of the genes express in a similar patterns between bat and human, except the low-density lipoprotein receptor (*LDLR*) gene, the receptor gene of human rhinovirus, which showes a low expression level in bat intestine and lung compared to that of human (Extended Data Fig. 1a). Therefore, at the level of bulk transcriptomic analysis, it is clear that the ability of bats to avoid overt disease from these viruses is not due to species expression of entry receptors in particular tissues.

**Figure 1.**
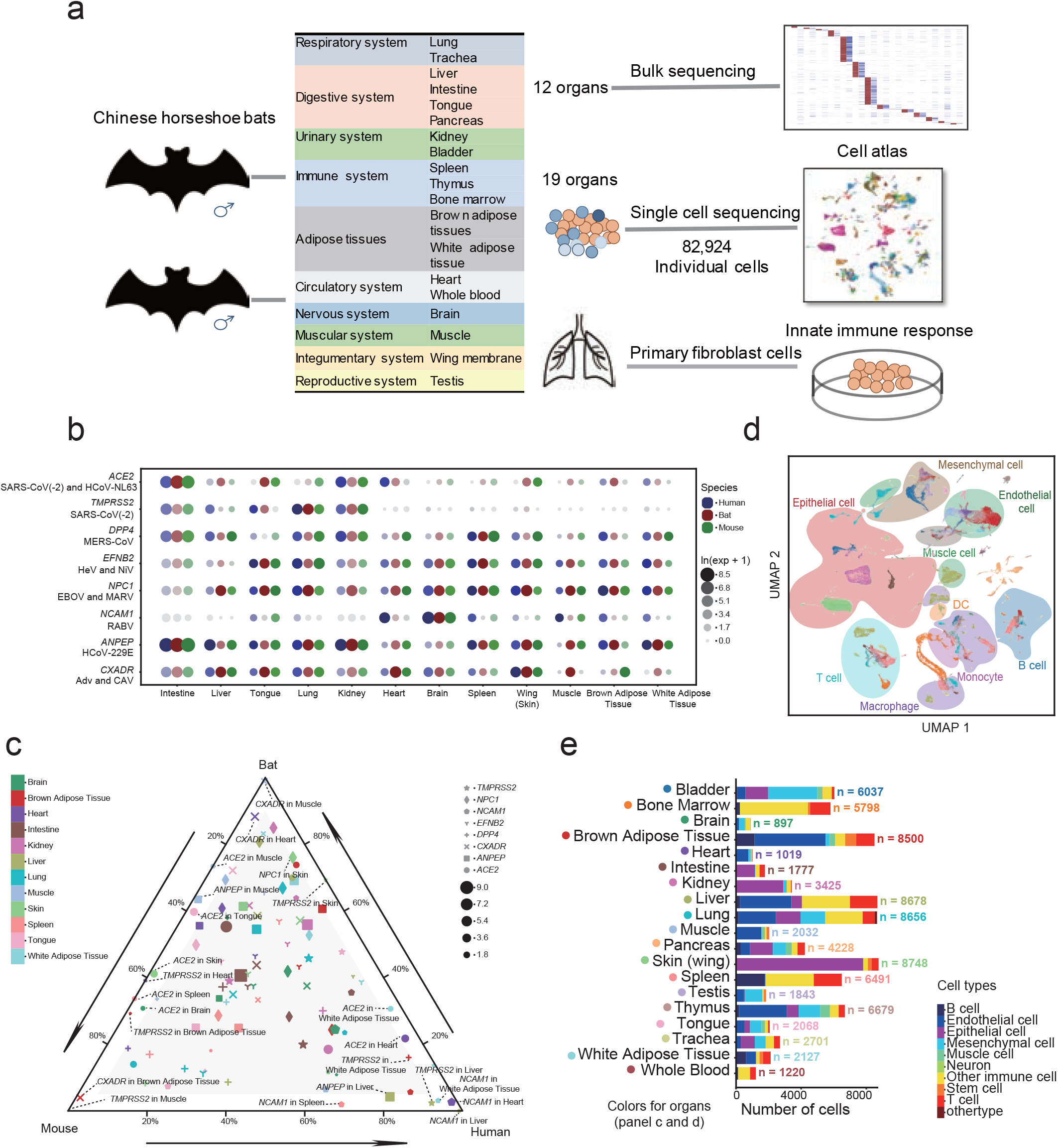
Overview of Chinese horseshoe bat cell atlas. a, Work flow of single cell sequencing. Cells from two male bat organs were processed for transcriptomic amplification, sequencing and data analyzing. b, The expression level of viral receptor genes across organs based on bulk-seq data between bat, human and mouse. The size of circle represents the gene expression level and the colors showed the species. c, Axis on the triangular representation of the distributions of viral receptor genes across the organs in bat, human and mouse. The size of the signals represents the mean gene expression showed as ln (expression + 1). d, UMAP plots of all cells, colored by organ, overlaid with the predominant cell type composing each cluster, n = 82,924 individual cells. e, The number of annotated cell types in each organ.

### Construction of single-cell atlas of bat

For sc-seq, nearly all tissues were obtained from both bats, with the exception of the intestine and white adipose tissue (Fig. 1a). Overall, 82,924 cells were retained after quality control. The median number of unique molecular identifiers (UMIs) per cell is 3,081 (Extended Data Fig. 1b). The organs were analyzed independently and cells were clustered according to the highly variable genes between cells by principal component analysis (PCA) and nearest-neighbour graph. A total of the 182 clusters were defined from the 19 organs (Extended Data Fig. 1c, Supplementary Table 1). The cell types in each cluster were annotated using known genes with differential expression between clusters. Significant differential transcriptional genes were observed across cell types, which encompass the gene module repertoire of the bat (Fig. 1d, 1e, Extended Data Fig. 1c). To ensure the accuracy of the single-cell (sc)-RNA seq data for cell typing, we further analyzed these differential transcriptional genes and found the similar expression pattern in corresponding organs in the bulk-seq data (Extended Data Fig. 1d). We constructed a bat cell atlas (http://bat.big.ac.cn/) for data accessing, enabling the searching of interested genes and browsing of single-cell data for all the organs.

To define whether there are varying gene transcription levels in different species at the single cell level, we explored differential gene expression by using the data from the bat lung and compared to that of human and mouse. In the bat lung, total 19 distinct clusters were classified, including 4 epithelial cells [alveolar epithelial type 1 (AT1) cell, alveolar epithelial type 2 (AT2) cell, ciliated cell, and mesothelial cell)], 3 endothelial cells (capillary type 1 cell, capillary type 2 cell, and lymphatic cell), 3 mesenchymal cells (adventitial fibroblast, alveolar fibroblast, and myofibroblast), 9 immune cell types (alveolar macrophage, interstitial macrophage, classical monocyte, non-classical monocyte, B cell, T cell, natural killer T cell, neutrophil and *ALOX5AP*^+^ macrophage) and one untyped cell cluster, which shows no specific expressed gene compared to other clusters (Fig. 2a, Extended Data Fig. 2 a-c). The differential genes expressed in epithelial cells, endothelial cells, mesenchymal and immune cells of lung across the species were then analyzed (Fig. 2b, Supplementary Table 2). In all the cell types, *DAZAP2*^21^, related to the regulation of innate immunity and *SUMO2*^22^ redundantly prevent host interferon response, were expressed at a higher level in bat compared to that of mouse. The genes related to cell proliferation (*MED28, TEMD3*) ^23,24^, cell cycle (*GATAD1*) ^25^, regulation of apoptosis and cell death (*ITM2C*) ^26^, host defense and inflammatory response (*CTSL*) ^27^expressed in higher levels in bat epithelial cells, endothelial cells and mesenchymal cells, while the *TAPBP*^28^, associated with antigen presentation, and *ARHGD1A*^29^, the regulator of Rho activity expressed higher in bat immune cells.

**Figure 2.**
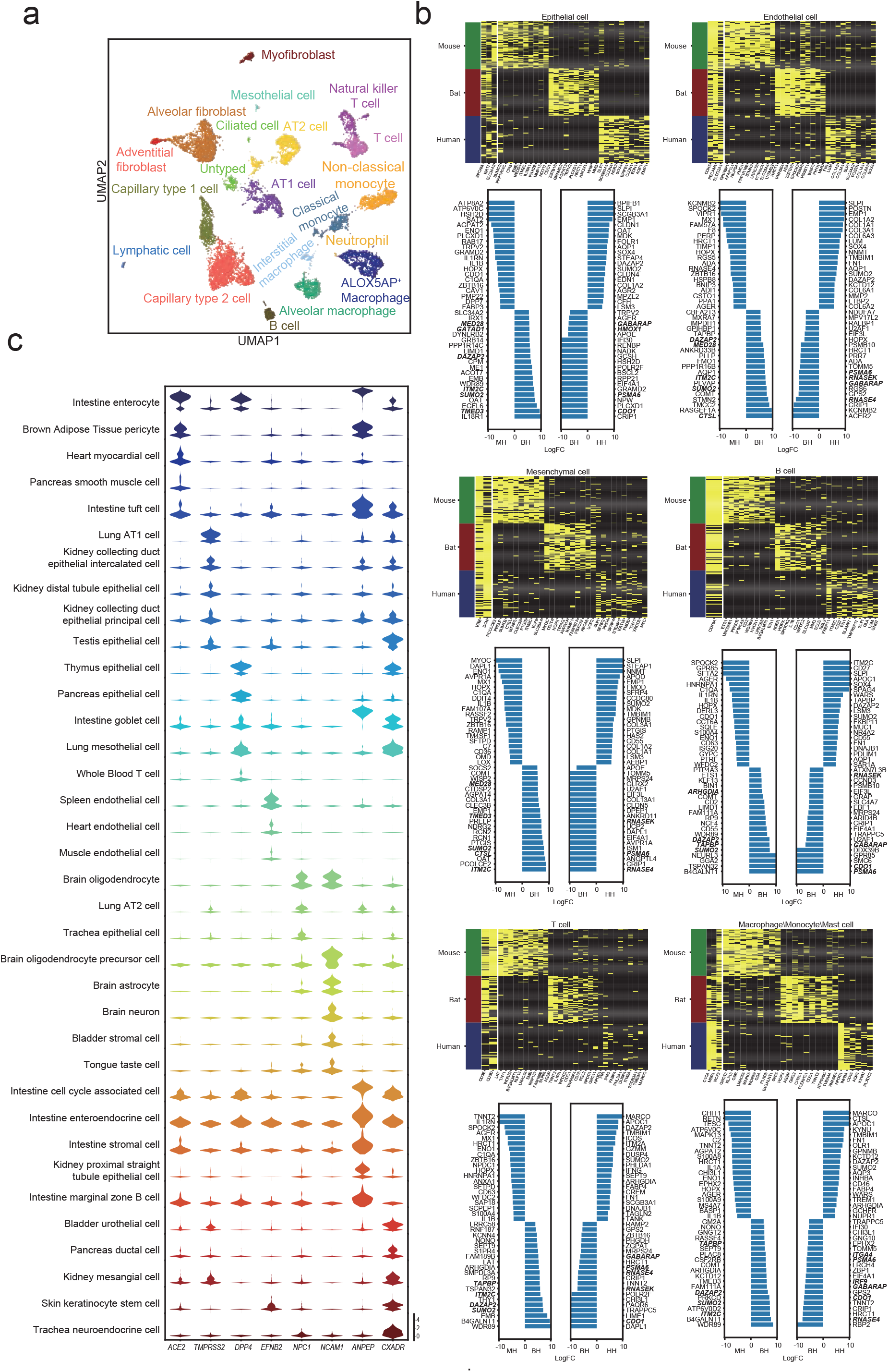
Differential expressions of bat lung cells compared to human and mouse, and the distribution of viral receptor genes across cell types. a, UMAP visualization and marker-based annotation of lung cells. Cells are colored by cell-type. b, Comparisons of the differential genes expressed in endothelial cells, epithelia cells, mesenchymal cells, and immune cells in bat compared to that of human and mouse. Differential expressed genes with p-adjust<=0.05 were analyzed. Results are visualized by heatmaps of normalized gene expression and histograms of fold change between cell types. c, Violin plots of viral receptor genes expression in top 5 cell types.

When compared to that of human, higher expression of several genes across the bat cell types were observed, including *PSMA6*, related to the inflammatory response^30^; *GABARAP,* a mediator of autophagy and apoptosis^31^; *CDO1,* the tumor suppressor genes ^32^; and *RNASE4*^33^, a member of RNase family associated with host defense-related activities, assumed to interact with pathogen-derived nucleic acid and facilitate their presentation to innate immune receptors within the cell as immunomodulatory proteins. Notably, the gene expressed ribonuclease kappa (RNASEK)^34^, recently identified as a host dispensable factor for the uptake of acid-dependent viruses, was highly expressed in bat lung cells (Fig. 2b and Supplementary Table 2). In bat lung epithelial cells, the gene encoded Heme oxygenase-1 (HMOX1), were observed at a higher level; this gene has been recognized as having anti-inflammatory properties and anti-viral activity^35,36^. In the bat monocytes, macrophages and mast cells, *ITGA4* and *IRF9* were expressed more highly compared to that of human. This well-characterized bat altas can gain an insight into cellular heterogeneity at the single cell resolution.

For viruses with respiratory and enteric tropism, we analyzed the viral receptor gene expression level across the cells (Fig. 2c, Extended Data Fig. 2d-g, Extended Data Fig. 3). This analysis shows that the respiratory virus receptor genes, *ACE2, DPP4, ANPEP* and *CXADR* are expressed at a high level in enterocytes, cell cycle-associated cells, enteroendocrine cells in the intestine, proximal straight tubule epithelial cells, and collecting duct epithelial cell (principal cells) in the kidney, and also in AT1, ciliated cells and mesothelial cells in the lung (Fig. 2c). *NCAM1* transcripts are mainly in oligodendrocyte precursor cells, oligodendrocytes, and astrocytes in the brain. *EFNB2* is mainly expressed in endothelial cells in the spleen, heart, and intestine, which is consistent with the NiV and HeV secondary replication sites, and corresponds to their important role in virus dissemination^19^. In addition, *EFNB2* is also expressed in intestine epithelial cells, but at a relatively low level, where NiV antigen have been identified in fatal human cases^19^. The expression of *NPC1* is broadly distributed in epithelial cells, endothelial cells, and mesenchymal cells.

The distribution patterns and expression levels of these receptor genes were then analyzed in human and mouse. We focused on the receptor genes (*ACE2, DPP4,* and *ANPEP*) of known zoonotic respiratory viruses, shared receptor of human and bat (*CXADR*) and *TMPRSS2* in the cell types in the trachea, lung, intestine and kidney. Bat exhibites more similar expression pattern to human in some organs (*ACE2* and *ANPEP4* in intestine epithelial cells; *TMPRSS2* in lung epithelial cells), comparing to that in mouse (Extended Data Fig. 2g). Although similar expression patterns were detected in bulk-seq data at organ level, the differences of cell types among species revealed by single cell data suggest bat as a better model for viral cross-species transmission research.

The other human viral receptor genes *NCL*, *CD55*, *HSPG2*, and *PDGFRA*, are expressed in the epithelial cells in both the respiratory tract and intestine, the viral tropism cell types (Extended Data Fig. 3). The transcripts of desmoglein 2 (DSG2), the receptor of Adv, Fc fragment of IgG receptor and transporter (FCGRT), fusion receptor of enterovirus B, are mainly expressed in epithelial cell and brush cell of the trachea, ciliated cell of the lung, and enterocytes in the intestine. These findings provide insights to understand the cellular tropism of respiratory tract and intestinal tract viruses (Extended Data Fig. 3).

This analysis of cell type specific gene expression data suggests that the distribution of viral entry receptor genes cannot explain the asymptomatic nature of viral infection in bats, and nor can it be explained by differential gene expression in those cell types. The molecular and cellular characteristics of the immune response in the bat were therefore then analyzed.

### The Adaptive Immune System: T and B cell clustering and analysis

Adaptive immunity in bats has been of great interest to understand their asymptomatic infection status as “viral reservoirs”. At the single-cell level, we analyzed the transcription features exhibited in immune cells. T cells differentially expressed CD3 genes in all organs are analyzed by using unsupervised clustering method implemented in Scanpy^37^. A total of 13 stable clusters are obtained, and each with unique signature genes (Extended Data Fig. 3a, 3b, Extended Data Fig. 4a-4c). In many organs, the number of activated T cells is much more than naïve T cells, such as liver, lung, trachea, intestine, pancreas, bladder, heart, kidney, and wing membrane (Extended Data Fig. 4d). Five clusters (C7, C8, C9, C11, and C12) express *CD8* genes, and seven clusters (C1‐C6, and C13) are composed of a mixture of CD4+ and CD8+ T cells. Most of C1, C3, C4 and C6 cluster are CD4+ T cell, while most of C2, C5, and C13 are CD8+ T cells. Cluster 10 is composed of CD3+CD4-CD8- T cells. Cells of C1_CD4T_N_ and C2_CD8T_N_ clusters expressing “naïve” marker genes such as *LEF1*, *CCR7*, and *TCF7* ^38^are mostly from spleen and bone marrow, respectively (Extended Data Fig. 4b, c, e). The cluster of C3_T_REG_-like is characterized by the expression of the *IL2RA* and *CCR8* genes, commonly associated with regulatory T cells (T_REG_-like). However, the *FOXP3* gene shows no expression in the clustered cells. The C4_T_CM_ cluster characterized by *CCR7*, *SELL*, and *GPR183* is composed of central memory T cells (T_CM_). The C5_T_EM_ cluster is closest to effector memory (T_EM_) T cells in many organs, in accordance with the expression of *CD44, CXCR3*, *GZMK*, *CCL5*, *CTSW* and *NKG7*, and the lack of expression of lymph node-homing receptors *CCR7* and *SELL*. The C6_T_EM_/T_H_1-like cluster characterized by *IFNG*, *CXCR3*, *GZMK*, and *CD4*, which mainly distributed in the intestine, lung, and liver. The C7_T_H_17 cluster contains T_H_17 cells mainly in the trachea and lung, with high level expression of *IL23R* and *RORC* genes. In addition to T_EM_ cells, recently activated effector memory or effector T cells (T_EMRA_) are also identified. The C10_T_EMRA_ differentially highly expressed effector molecules such as *NKG7*, *ZNF683*, *CTSW*, *CCL5*, *GZMA*, *XCL1*, *KLRG1,* and *TBX21*. It has been reported that chemokines *XCL1* and *CCL5* derived from NK cells recruit cDCs into the tumor microenvironment, which are critical for antitumor immunity^39^, while KLRG1+TBX21+ T cells are long-lived effector cells, which contribute to infection control (Fig. 3b, Extended Data Fig. 4a-c).

**Figure 3.**
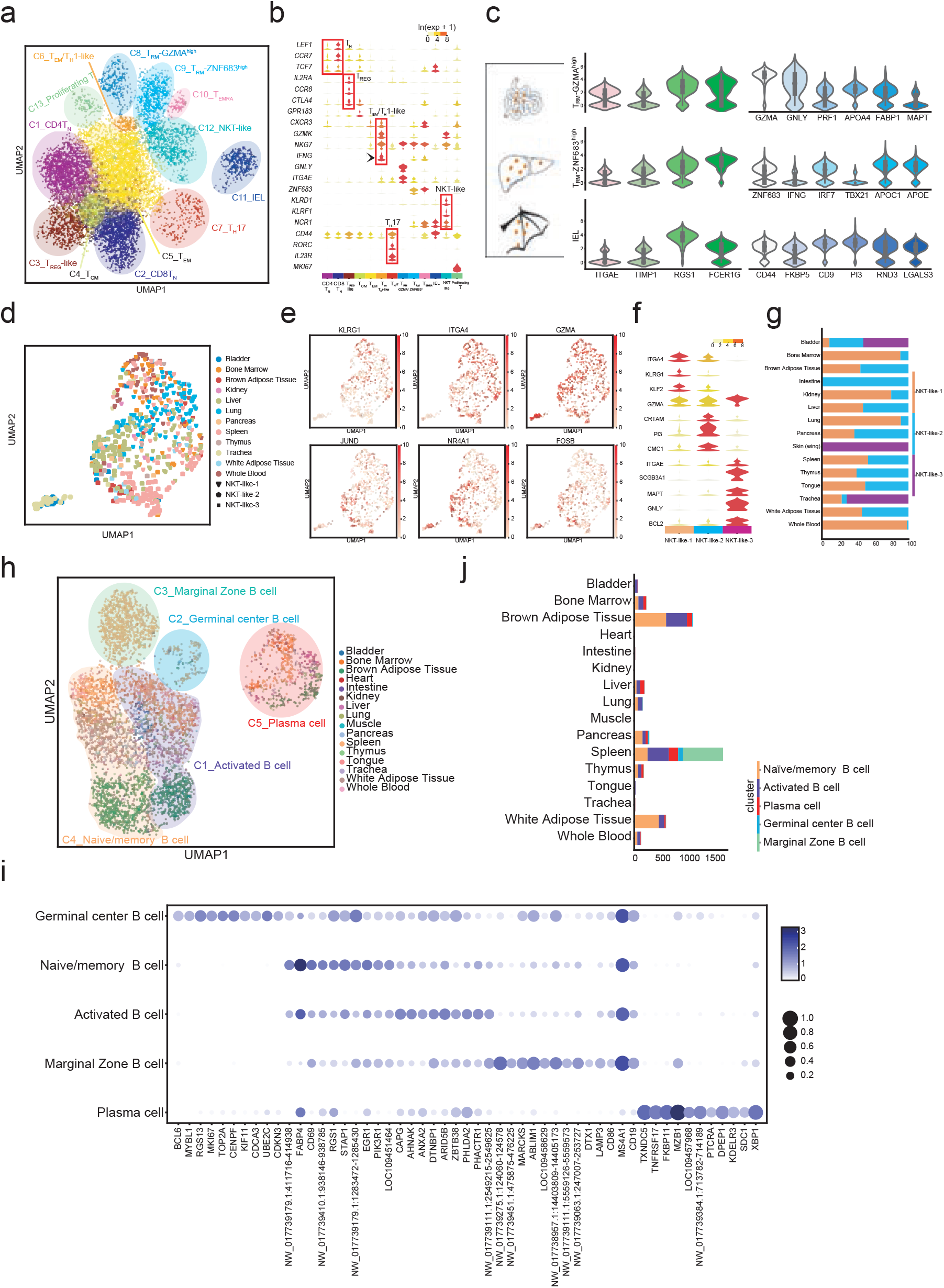
Analysis of bat T and B cells. a, UMAP visualization of all T cells from Chinese horseshoe bats (*Rhinolophus sinicus*), showing the formation of 13 main clusters shown in different colors. The functional description of each cluster is determined by the gene expression characteristics of each cluster. n = 9,663 individual cells. b, Violin plots showing the enriched transcripts of different T cell clusters. c, Violin plots showing the enriched transcripts of T_RM_-GZMA^high^, T_RM_-ZNF683^high^, and IEL. d, UMAP visualization of NKT-like-1, NKT-like-2, and NKT-like-3 cells. Colors indicate different organs, and shapes indicate cell types. e, UMAP plots showing expression of selected long-lived effector genes and immediate early genes (IEGs) in this dataset. f, Violin plots showing the enriched transcripts of NKT-like-1, NKT-like-2, and NKT-like-3 cells. g, Tissue preference of each NKT-like cell cluster estimated by proportion. h, UMAP visualization of B cells from Chinese horseshoe bats (*Rhinolophus sinicus*), showing the formation of 10 main clusters shown in different organs. The functional description of each cluster is determined by the differential expressed genes. i, Dot plot visualization of selected marker gene for each cell type. The size of the dot encodes the percentage of cells within a cell type in which that marker gene was detected, and the color encodes the average expression level. j, Distribution of B cell types in different organs.

The cells in C8_ tissue-resident memory T (T_RM_)-GZMA^high^, C9-T_RM_-ZNF683^high^, and C11_IEL clusters express the *ITGAE* gene, a known marker of T_RM_ cells. They share signature genes, such as *TIMP1*, *RGS1*, and *FCER1G*. The cells in C8_T_RM_-GZMA^high^ are predominantly from the intestine and express cytotoxic molecules such as *GZMA*, *GNLY*, *PRF1,* and *CCL5*. Most of the cells in C9_T_RM_-ZNF683^high^ are from the liver, which express higher levels of effector molecules such as *ZNF683, NKG7, XCL1* and *CCL5*, and interferon stimulating genes (ISGs), including *IFNG* and *IRF7*. The cells in the C11_IEL cluster are detected exclusively in the wing membrane, which are considered as intraepithelial lymphocytes (IEL) as they highly expressed natural killer cell receptor genes, *NCR1* and *KLRB1*^40^. These cells also display overwhelmingly active molecules *CD44*^41^, chemokine *XCL1*, and ISG *CD9* (Fig. 3c, Extended Data Fig. 4e).

The C12_NKT-like cluster characterizes NK cell receptor genes, including *KLRB1*, *KLRD1*, *KLRF1*, *KLRG1*, *NCR1*, and *NCR3* genes. All the genes express overlapping with the *CD3* gene and the cluster is considered as NKT-like cells. The C12_ NKT-like cluster is composed of three subsets and each subset expressed distinct high level genes in different organs (Fig. 3d-3g). The subset-1 which highly express *KLRG1, GZMA*, and *ITGA4* is considered as long-lived effector NKT and contribute extensively to immune surveillance^42^. The subset-2 highly expresses active gene *CRTAM* and *CD69*, peptidase inhibitor gene (*IP3*), chemokine *XCL1*, and immediate early genes (IEGs), such as *JUND*, *NR4A1*, and *FOSB*. IEGs are rapidly activated at the transcriptional level in the first round of response to stimulation prior to any nascent protein synthesis. These data suggest subset-2 NKT-like cells are in active states. The subset-3 NKT-like cells highly express *SCGB3A1*, *SCGB3A2, MAPT*, *GNLY*, and *BCL2*. The *SCGB3A1* is a tumor suppressor gene, while *SCGB3A2* is a negative inflammation response gene. C13_proliferating T cells are significantly enriched in the expression of cell cycle genes (Fig. 3f, Extended Data Fig. 4a-4c), such as *MKI67*, *UBE2C*, *CENPF*, *PCLAF*, *TOP2A*, indicating the proliferative states of the C13 cells.

B cells are annotated into five clusters according to the marker genes (Fig. 3h, 3i). C1 cluster contains activated B cells expressing high levels of *CD86*, *CAPG*, *AHNAK*, *ANXA2*, and *PHACTR*1. C2 is defined as germinal center B (GC B) cell with the specific expression of BCL-6, and cell cycle-related genes (*MKI67*, *TOP2A*, *CENPF*, *CDCA3*, and *CDKN3*) C3 cluster is defined as marginal zone B (MZB) cell, which highly expressed *MZB1*, *DTX1*, and NW_017739275.1:124060-124578 (corresponding to *Rousettus aegyptiacus* complement component 3d receptor 2 (CR2) gene). Some MZB cells express *SDC1*, indicating a conserved maturing location of plasma cells. The C4 cluster is characterized by higher expression of *FABP4*, *RGS1*, *STAP1*, and *PIK3R1*, which are defined as naïve/memory B cells. The C5 cluster is annotated as long-lived plasma cells, in which the transcription genes *SDC1*, *PRDM1*, and *XBP1* and marker genes *SDC1* and *TNFRSF17* expressed at a high level, while *CD19* and *MS4A1* were under-expressed. A majority of B cells were in the spleen and adipose tissues (Fig. 3j).

### The gene expression patterns of mononuclear phagocytes

Mononuclear phagocytes (MNPs) play a critical role in pathogen sensing, phagocytosis, and antigen presentation. The MNPs in bat tissues are profiled and 12 clusters are grouped according to differentially expressed transcripts (Extended Data Fig. 5a, Supplementary Table 1). C9_granulocyte-monocyte progenitor (GMP) is identified in bone marrow, with specific expression of *CTSG*, *MPO*, *ELANE*, and *RTN3*. Two clusters of monocytes are identified, including C3-classical monocyte (cMo) and C1_nonclassical monocyte (ncMo). Six clusters are macrophages, including C0_CSF3R^high^ macrophage (CSF3R^high^ Mac), C10_CD300E^high^ macrophage (CD300E^high^ Mac), C2_Kuffer cells (KC), C4_SPIC^high^ macrophage (SPIC^high^ Mac), C5_LYVE1^high^ macrophage (LYVE1^high^ Mac), and C6_CCL26^high^ macrophage (CCL26^high^ Mac). The macrophage clusters are characterized by unique expressed genes, such as CSF3R and *SERPINA12* in CSF3R^high^ Mac, *CD300E* and *LPO* in CD300E^high^ Mac, *MARCO* and *CLEC4G* in KC, *CNTNAP2*, *SPIC*, and *VCAM1* in SPIC^high^ Mac, *LYVE1* and *DAB2* in LYVE1^high^ Mac, *CCL26* and *SCD* in CCL26^high^ Mac (Extended Data Fig. 5b, Supplementary Table 1). KC, SPIC^high^ Mac, LYVE1^high^ Mac, and CCL26^high^ Mac are tissue-resident macrophages with high expression of *C1QA, C1QB, C1QC*, and *MAFB*. A total of about 600 differentially expressed genes are identified in cross different tissue-resident macrophage populations. The differential expressed genes in CSF3R^high^ Mac, mainly identified in the intestine, are enriched in leukocyte migration, leukocyte chemotaxis, and positive regulation of response to external stimulus in GO annotations (Extended Data Fig. 5c and 5d). SPIC^high^ Mac mainly comes from the spleen. The enriched genes for GO annotations are mainly responsible for signal transduction, activation of the immune response, and regulation of monocyte chemotaxis. LYVE1^high^ Mac contains many organs macrophages, such as the bladder, pancreas, thymus, adipose tissue, heart, tongue, trachea, testis, kidney, lung, muscle, and intestine. The enriched gene functions mainly include wound healing, cell migration, and protein activation cascade. CCL26^high^ Mac cluster contain macrophages of the lung and trachea, which are mainly associated with lipid transport and metabolic process, phagocytosis, and regulation of endocytosis (Extended Data Fig. 5c, d).

Four DCs subtypes are determined (Extended Data Fig. 5a). C7_cDC1 is characterized by *FLT3*, *CLEC9A*, *XCR1*, *IRF8*, and *CPVL*. C12_pDCs highly express *TCF4*, *IRF8*, *IL3RA*, *IRF4*, *LAMP3*, *BCAS4*, and *GPM6B.* C11 is annotated as activated DCs, with high expression of DC hallmark receptor gene *FLT3*, activation marker gene *LAMP3*, co-stimulatory molecule genes *ICOSLG* and *CD83*, and chemokine receptor genes *CCR7* and *IL7R*. C8_Langerhans cells (LCs) are mainly located in the wing membranes, with highly expressed genes of *FLT3*, *RUNX3*, *EPCAM*, and *TACSTD2* (Extended Data Fig. 5b, c). Collectively, monocytes, macrophages, and DCs display distinct gene landscapes, which likely form the basis of MNPs specificity and plasticity. The distinct gene profiles of MNPs may contribute to the critical role of MNPs in pathogen sensing, phagocytosis, antigen presentation, tissue function and homeostasis.

### Innate immunity: the response of bat primary lung fibroblast cells against RNA virus infection

We then studied the innate immunity activities since it is the first-line to control the virus infections. Real-time quantitative PCR analysis revealed that innate immunity related genes, such as retinoic acid-inducible gene-I (*RIG-I*), melanoma differentiation-associated protein 5 (*MDA5*), toll-like receptor (*TLR*) 3, *TLR*7-9, interferon regulatory factor (*IRF*) 3, *IRF*7, *IFN*α, β, ω, and γ, are expressed in various tissues (Extended Data Fig. 6). All of these genes are highly expressed in the spleen and white adipose tissue. Furthermore, *TLR*3 and *TLR*7 are highly expressed in the intestine. *RIG-I*, *MDA-5*, *TLR-3*, *TLR-8* and IRF3 are highly expressed in the lung (Extended Data Fig. 6).

To analyze the innate immune activities at the lung cellular level, we isolated primary bat lung fibroblasts (BPFs). According to the transcriptomic data, the lung stomal cells constitutively expressed innate immune genes (Extended Data Fig. 7). An RNA virus, vesicular stomatitis virus (VSV), and/or the analogs stimulating the signaling pathways, were used to treat BPFs and human primary lung fibroblasts (HPFs) (Fig. 4a). The cells were stimulated with poly (I:C), R848, the analogs of RIG-I/MDA-5, TLR3, and TLR7/8, respectively, as well as VSV. At 4 hours (h), 8h, 12h and 24 h after treatment, the expression levels of RIG-I, MDA5, IFNα, β, IL-6 and TNFα were analyzed. In HPF, the transfection of poly (I:C) induces the expression of IFNα, β about 4,000-fold compared to untreated cells at 4h post-treatment, while the incubation of poly (I:C), for the purpose of stimulating the TLR3 pathway showed similar results (Fig. 4c-4e). However, the extent of IFN-β mRNA induction was much lower in BPFs when compared with HPFs after 4h post-treatments (*p*=0.000, student *t* test). The transfection of R848 induces higher expression levels of IL-6 and TNFα mRNA in HPFs, but not in BPFs (Fig. 4b). Similar results were obtained in VSV-infected cells (Fig. 4f). Although the VSV replicates in a low level in BPFs compared to that of HPFs, the mRNA levels of MDA-5 and RIG-I increased slightly. However, the transcription of IFN-β, IL-6, and TNF-α does not increased significantly compared with that in HPFs.

**Figure 4.**
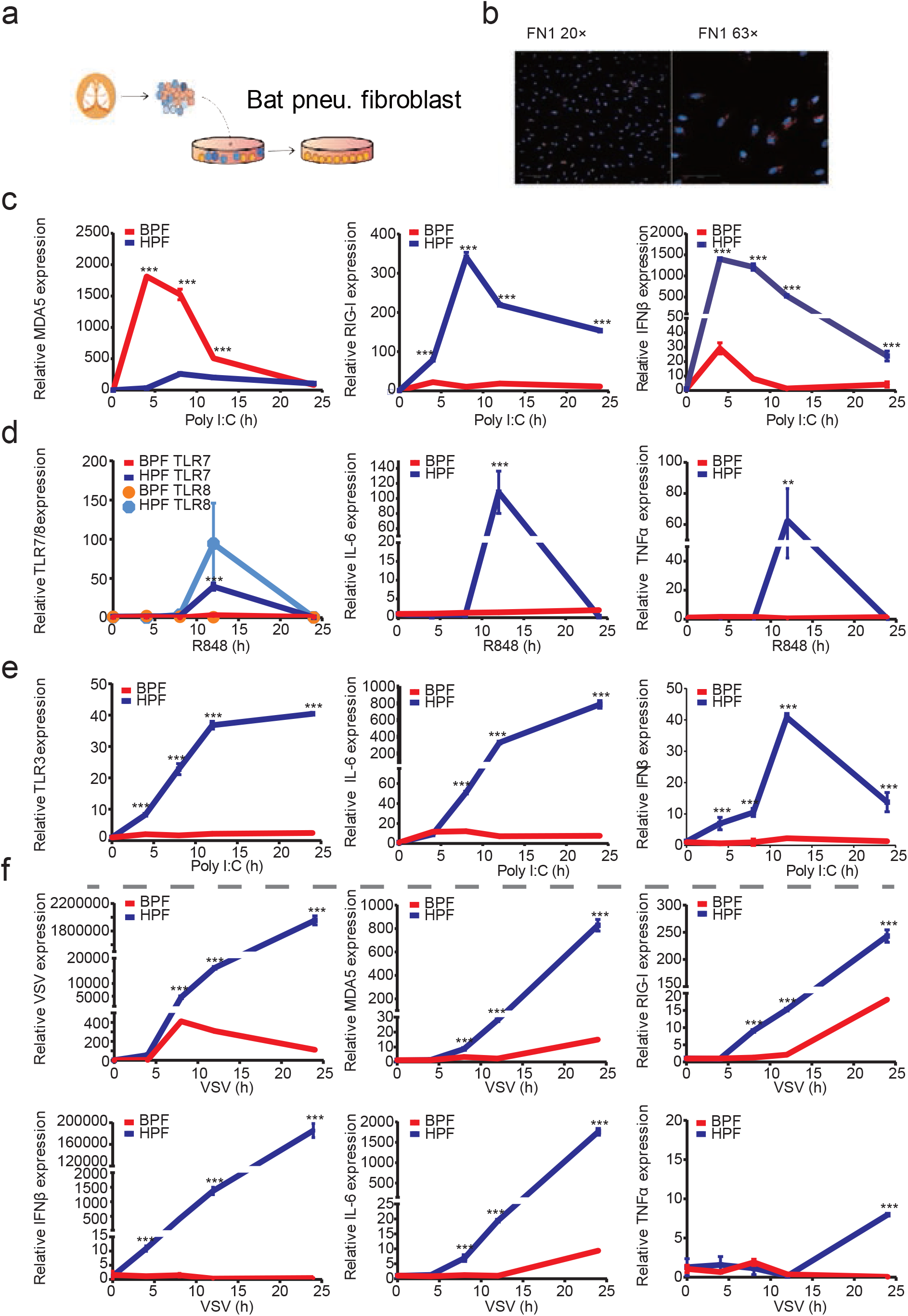
Expression levels of type I interferon in bat primary lung fibroblast. a, Work flow of bat pulmonary primary fibroblast isolation. b, Cell type confirmation by single-molecule fluorescence in situ hybridization with fibronectin 1 RNA probe. c, Expressions levels of MDA5, RIG-I, IFNβ after the transfection of poly (I:C). d, Expressions levels of IL-6, TNFα after the transfection of B848. e, Expressions levels of IL-6 and IFNβ after the inoculations of poly (I:C). f, Expressions levels of MDA5, RIG-I, IFNβ, IL-6, and TNFα after the infection of vesicular stomatitis virus (VSV). Error bars represent standard deviation.

## Discussion

The single-cell transcriptome data obtained from the model organism mouse and human has established a reference database in mammals for deep molecular annotation of cell types^13–15^. Here, we have created a parallel cell atlas in an important organism trapped from the wild.

Most emerging infectious diseases in humans are dominated by zoonoses^43^. Over millions of years, bats have evolved a special ability to carry a variety of viruses but show little or no signs of disease. However, many of these viruses result in devastating infection when they cross the species barrier to humans^44,45^. Bats are social animals and diverse viruses circulate within the colony, allowing them to be important natural viral reservoirs^45^. Viral strains or mutatants adapted to human beings or other species may occur during the circulation, which can spill over to human beings or other animal species (intermediate species). This may result in epidemics or outbreaks in human beings by bat-human or bat-intermediate hosts-human transmissions^46^. Therefore, bat-borne viruses etiologically play a pivotal role in human emerging infectious diseases (EIDs), and understandings in how bat carry and transmit viruses has become a priority issue for the EIDs alert and prevention^47,48^. The infection and replication of viruses rely on specific host cells due to their parasitic nature. Interactions between the virus and the host affect the infections and the replicate abilities of the virus, and we have lacked an understanding of how host cell biology determines whether the infection is asymptomatic or pathogenic. Furthermore, the pathogen’s shedding route is highly dependent on the replication tissue sites. However, the lack of precise annotation of cellular composition of bat organs/tissues has hindered the understanding on the many key aspects of zoonotic virus origin in bats, e.g. the mechanisms of asymptomatic bearing of diverse viruses, tropism and tissue targeting, virus shedding route and interspecies transmission, etc. It is critically needed to elucidate the cellular composition as well as their functions and interplay in various bat organs/tissues. However, it is still hard to create bat cell atlas by conventional immunological or histological approaches due to the lack of sufficient antibody reagents at present. Single-cell sequencing does not rely on cell surface protein markers and antibodies. By clustering based on the transcription characteristics and specifically transcribed genes, cells in organ/tissues can be classified by sc-seq. In this study, we developed the first bat cell atlas by single-cell RNA-seq, which provides a powerful tool for in-depth understanding of the cellular mechanisms by which how bats carry, shed and cross-species transmit viruses.

Binding and entry into the host cell is the first step of virus infection. The specific receptor molecules on host cell membranes govern whether a virus can enter and infect the cells. The bat-borne viruses, such as SARS-CoV, MERS-CoV, HeV, NiV, RABV, EBOV, and MARV, have emerged for more than twenty years and resulted in human infections in the world. A crucial factor for these viruses infected human is that these viruses could enter the human cells through specific receptors. In this study, it was found that bat-borne viruses share similar receptor distributions and expression in bat and human. For example, the the NPC1 (receptor of EBOV and MARV) are distributed in most organs in bat and human, with similar expression levels. It has been reported that patients succumbed to EBOV and MARV infections have extensive necrosis in parenchymal cells of many organs, including liver, spleen, kidney, and gonads^49^, which is consistent with the distribution of receptors in bat in this study. Further, EBOV and MARV have a broad cell tropism. It has been verified from fatal human cases or experimentally infected nonhuman primates that EBOV and MARV could infect and replicate in monocytes, macrophages, dendritic cells, endothelial cells, fibroblasts, hepatocytes, and several types of epithelial cells^20,50^. The distributions of NPC1 in immune cells and nonimmune cells further support the broad cell tropism of EBOV and MARV. ACE2, the SARS-CoV and SARS-CoV-2 receptor, is expressed in epithelial cells of the lung and the small intestine, which are the primary targets of the two CoVs, as well as in the heart, kidney, and other tissues^16,51^. In this bat data, it was found that *ACE2* is mainly expressed in different epithelial cells of the intestine, lung and trachea, such as enterocytes, enteroendocrine cells, goblet cells and tuft cells, and lung and trachea ciliated cells. *TMPRSS2* showes higher expression in enterocyte of the intestine, AT1 in the lung and collecting duct cells in the kidney. However, most of the receptor genes of respiratory viruses show a higher expression level in bat intestine epithelial cells. Previous studies also showed that SL-CoVs are majorly detected in the intestine of bats^3,4^. These findings indicate that intestine is probably a major site where many viruses reside and replicate, such as SL-CoVs^4,7^. This may facilitate their dispersal in the nature as feces harboring the shed viruses can touch other animal species more effectively than the respiratory route. These data suggest that the similar receptor distribution patterns between bat and human may be one of the bases of cross-species spread of bat borne viruses. Further, some virus receptors, such as SARS-CoV, SARS-CoV-2, MERS-CoV, HeV, NiV, RABV, EBOV, and MARV, are also co-located in the lung, bladder, and intestine of both species, which is critical in virus transmission. The clarifications on the consistence and difference of surface molecular patterns between bat and human across cell types, for example, the viral receptor gene, would be informative to assess the cross-species transmission risk of bat borne viruses.

The knowledge of cellular immunity of bats is quite limited. According to the sc-seq data, we found that CD8^+^ T cells were predominant over CD4^+^ T cell in Chinese horseshoe bats (*Rhinolophus sinicus*). All tissue-resident memory T cells, including C8-T_RM_-GZMA^high^, C9-T_RM_-ZNF683^high^, and C11_IEL, are CD8^+^ T cells. These T_RM_ cells highly expressed many effector molecules such as *GZMA*, *GNLY*, *PRF1*, *ZNF683*, *NKG7*, *XCL1* and *CCL5*, and so forth. Microbes most often attack body surfaces and mucosal sites. T_RM_ cells lie in frontline sites of infection and need not proliferate. They are anatomically positioned to respond most immediately, which contribute to pathogen control after the initial infection. In addition to T_RM_ cells, most of T_EM_ and NKT-like cells are CD8^+^ T cells, and they express many effector molecules and IEGs. Specifically, subset-2 NKT-like cells highly expressed IEGs, which can induce a rapid response to stimuli before new protein synthesis. At the same time, *CD69*, an activation inducer molecule, displayes high enrichment in subset-2 NKT-like cells. These data imply an active state of the NKT-like cells in subset-2. Collectively, these predominant CD8+ T cell and their functional states suggest that the immune baseline level of the bat is quite high and is geared towards fighting microbe infections. The resident tissue CD8+T cells and NKT-like cells with the higher expression level of immediate early genes indicates effective cellular immunity response restricting viral infections in bats.

It has been reported that *KLRG1*^+^ T cells and NKT are long-lived effectors and optimally provide immediate protective immunity against certain pathogens^42,52^. In this study, subset-1 NKT-like cells display high expression of *KLRG1*, with co-expressed *ITGA4* and *GZMA*. These NKT-like cells distribute in different tissues of the bat. This will help the bat to better conduct immune surveillance and fight against infections and tumors. T_EMRA_ cells present high expression of *KLRG1* and *TBX21* in the bat, which have been reported to be expanded and maintained long term following boosting, without losing their protective superiority^52^. T_RM_ cells resident in the local environment long after peripheral infections subside. If an infection is localized to peripheral or extralymphoid compartments, T_RM_ cells would provide superior immune protection than circulating memory T cells^53^. The circulating CD8+ memory T cells is failed to control the wing membrane infection with HSV, while the T_RM_ cells in the wing membrane provide local protection against infection in the absence of ongoing T-cell stimulation. We found that there are many T_RM_ cells in the bat intestine, liver, and wing membrane, indicating an activated adaptive immunity, which may offer effective barrier immune protection for bat.

The highly activated cellular immunity may protect bats from viral damage. But how can the virus reside and replicate in bats? The tolerance of viral infections in bats appears to involve a balance between viral clearance and host tissue damage promoted by proinflammatory effectors. The innate immune system is the first defense against invading pathogens in mammals and type I IFNs are induced very early in viral infection. The magnitude and nature of the IFN response determines whether the resulting effects on the host are harmful or beneficial^54^. In Chinese horseshoe bats (*Rhinolophus sinicus*), the IFN gene loci are still not clarified clearly. However, the critical components of the IFN signaling pathway have been investigated^55,56^. For example, the sequences of RIG-I, STAT-1 and IFN-β have close homology with human, mouse, pig and rhesus monkey in immortalized embryonic fibroblast (BEF) cell lines from *Rhinolophus affinis* and *Rhinolophus sinicus*^57^. Our data show that the critical host genes in the IFN signaling pathways are expressed across the cell types. To characterize the innate immunity in bat, we isolated BPFs from one of the Chinese horseshoe bat (*Rhinolophus sinicus*) lungs. We found that the induction of most major pathogen associated recognition pattern (PAMP) receptors, including RIG-I, TLR-3, and TLR7/8 as well as IFN-β and proinflammatory factors, such as IL-6 and TNFα, are very low in BPFs compared to HPFs when stimulated by polyI:C and VSV. These data indicated that the signaling pathways of innate immunity in bat are tightly suppressed. The low level innate immune response may enable the bats to asymptomatically harbor viruses.

In conclusion, we show here a first comprehensive bat cell atlas based on single-cell transcriptional landscape of 19 organs from Chinese horseshoe bat. By combining the sc-seq and bulk-seq data, we characterized the distribution patterns of multiple known human viral receptors in bat and human across organs and cell types. We also demonstrate an orchestration of highly activated adaptive immunity and suppressed innate immunity status, which may form a precise immune hemostasis which allow the virus harbor in bats without pathological damage. Our findings provide insights into the cellular mechanisms to enable bats to serve as natural viral reservoirs, largely informing an active alert and control of epidemics caused by bat borne viruses.

### Online content

The methods, additional references, source data, statements of data availability and associated accession codes are available online.

## Methods

### Bat, organs and single cell preparation

The two male Chinese horseshoe bat (*Rhinolophus sinicus*) were obtained in October, 2018 from Anhui province, China. The bats were placed separately and transferred to the lab. The species of each bat was identified by field biologists and recorded. After anaesthetization with pentobarbital sodium (75mg/kg), bats blood was drawn via cardiac puncture using sterile syringes, then the other organs and tissues were isolated as followed, pancreas, intestine, spleen, liver, kidney, brown adipose tissue (interscapular adipose tissue), white adipose tissue (visceral and subcutaneous adipose tissue), thymus, heart, lung, trachea, bladder, testis, tongue, brain, muscle, wing membrane, and bone marrow (forelimb bones). The cell suspensions from each tissue and organ were prepared and the details were available as followed. The experiments and programs were reviewed and approved by the Institutional Animal Care and Use Committee of the Institute of Laboratory Animal Science, Peking Union Medical College (BYS18003).

### Single cell preparations

The whole blood was quickly transferred into 1.5ml sterile tubes with anticoagulant EDTA and mixed gently, then suspended with 0.5 ml of red blood cell lysis buffer. Cell suspension was incubated on ice for 1 min and lysis reaction was quenched by adding 10 ml Dulbecco’s Phosphate Buffered Saline (DPBS) with 2 mM EDTA and 0.5% BSA. Cells were collected at 200g× for 5 min at 4 □ and washed with DPBS for two times to remove the lysis buffer. The viability of cells was determined with trypan blue stain method by calculating the rate of bright cells (viable) to stained cells (no-viable) with hemocytometer.

Spleens were rinsed and dissected quickly in cold DPBS, then squeezed to pass through a 70 μm strainer using plungers. Cells were collected into a 50ml centrifuge tube, and then centrifuged at 300 g× for 5 min at 4□. Cells were resuspended with 3 mL of red blood cell lysis buffer. Cell suspension was incubated on ice for 1 min and lysis reaction was quenched by adding 20 ml DPBS with 2 mM EDTA and 0.5%BSA. Cells were collected at 300g× for 5 min at 4□ and washed for 2 times with DPBS, then counted with hemocytometer after Trypan blue staining as described above.

Bone marrow was isolated from bat forelimb bones. Both ends of bones were carefully trimmed to expose the interior marrow shaft after removed the wing membrane and muscles. The bone marrow cells were flushed by using 1 ml syringe with DPBS for several times, all the cells were collected into a 70 μm strainer hanging on a 50 ml centrifuge tube. Bone marrow cells on the strainer were gently squeezed to pass through by using plungers. Cells were centrifuged at 200 g for 5 min at 4□ and resuspended with red blood cell lysis buffer. The red blood cell lysis, washing, and cell counting process were similar as above.

Other organs were minced into pieces on ice with sterile scissors respectively. Tissue pieces were respectively transferred into a 15 ml centrifuge tube and suspended with 5 ml of enzymatic digestion buffer. Samples were treated with different enzymes formula. The bladder, brain, brown and white adipose tissue, intestine, liver, lung, pancreas, testis, thymus, and trachea were respectively digested in enzymatic digestion buffer with 0.4mg/ml collagenase IV, 0.4mg/ml collagenase/dispase, 30U/ml DNase, 0.5% BSA in HBSS, at 37□, 100rpm for 30min. Heart was digested with 1mg/ml collagenase/dispase, 30U/ml DNase, 0.5% BSA in HBSS at 37□, 100rpm for 45min. The intestine, muscle and testis were digested in enzymatic digestion buffer with 0.4mg/ml collagenase II, 30U/ml DNase,0.5% BSA in HBSS, at 37□, 100rpm for 60min. The kidneys and wing membrane were digested with 0.25% Trypsin and 30U/ml DNase, at 37□ for 15min and 30min, respectively. Tongue was digested with 0.4mg/ml collagenase IV, 30U/ml DNase, 0.5% BSA in HBSS, at 37□, 100rpm for 60min. Tissue pieces were pipetted up and down gently for several times to dissociate into single cells during digestion. After passing through a 70 mm strainer, the dissociated cells were centrifuged at 300 g for 5 min at 4□ and treated with red blood cell lysis procedure. All the treated cells were finally diluted to a density of 1000 cells/μl in DPBS with 0.4% BSA.

### Single-cell sequencing library construction and sequencing

Sc-seq libraries were constructed by using the Single-Cell Instrument (10 ×Genomics, Pleasanton, CA) with Chromium v2 single cell 3’ library and gel bead kit V2 (10 × Genomics, Pleasanton, CA). In brief, cell suspensions were diluted in DPBS with 0.04% BSA to concentration of 1,000 cells/μl and the concentration was measured with haemocytometer. The volume of single cell suspension required to generate 4,000 single cell gel beads in emulsion (GEMs) was loaded into a separate channel on the Single Cell 3′ Chip. The final libraries were qualified with Agilent 2100 Bioanalyzer (Agilent Technologies, Santa Clara, CA, USA), and quantified by qPCR with the quantification kit (Tian Gen company, China) for Illumina with QuantStudio 12K Flex Real-time PCR system (Thermo Fisher Scientific, Waltham, MA, USA). Libraries were diluted to 2 nM in each and pooled with equal volume before sequenced on Hiseq X Ten (Illumina, Inc., San Diego, CA, USA) with 150-bp pair-end strategies.

### Bulk RNA-seq library construction and sequencing

Approximately 30-50 mg of each tissue was collected from each bat and homogenized using FastPrep-24 system (MP Biomedicals, France) in 1ml of TRIzol (Invitrogen, Carlsbad, CA). RNA was then extracted following standard protocol. The RNA qualities determined by RNA Integrity Number (RIN) were assessed on an Agilent Bioanalyzer RNA 6000 nano chip (Agilent Technologies, Santa Clara, CA, USA). The libraries were constructed by using the NEBNext ultra II RNA library prep kit (New England BioLabs, Ltd., USA) and the qualities were analyzed with Agilent 2100 Bioanalyzer (Agilent, Santa Clara, CA). Libraries were then sequenced on HiSeq X Ten (Illumina) with 150-bp pair-end strategies.

### Single cell sequencing data processing and clustering

Sequencing reads were first aligned to Rhinolophus sinicus genome (GCA_001888835.1) using CellRanger (version 3.0.0, 10× Genomics) with default parameters. Then the sequencing data was processed for filtering, variable gene selection, dimensionality reduction, and clustering by using Scanpy package (version 1.3.7). Cells with fewer than 500 detected genes were excluded, as well as expressed fewer than 1,000 unique molecular identifiers (UMIs). Gene expressions were normalized as divided by total UMIs of each cell and multiplied by 10,000. Highly variable genes were selected by coefficient of variation with cutoff of 0.5. After log-normalized and scale, the data dimensionality was reduced by principal component analysis (PCA) by variable genes. Neighborhood graph of observations were computed based on the Euclidean distance and parameters were adjusted for each tissue. The Batch balanced KNN package (bbknn, version 1.3.1) was used for batch correcting followed the procedure by identifying the K nearest neighbours for each individual cell. Cluster cells using the Leiden algorithm and cell type of each cluster was determined by using the abundance of known marker genes. Cells were visualized using UMAP method which is a manifold learning technique suitable for visualizing high-dimensional data. To compare the human and mouse gene expression with the bat respectively, we use the public databases of human metadata available on GEO (accession GSE130148) and mouse metadata online (http://tabula-muris.ds.czbiohub.org/) for later normalization and gene expression comparison.

### Bulk sequencing data processing

The gene expression profiles of bat tissues from bulk-seq data were performed following typical RNA-Seq procedure with reference genome. The raw-reads were treated to generate clean-read datasets by the following procedure. Reads with adaptors or containing unknown nucleotides more than 5% were removed directly. The low-quality reads containing more than 20% suspect-nucleotides of Phred Quality Score less than 10 were then filtered out. The qualified reads were evaluated to trim unreliable ends containing more than 3 successive suspect-nucleotides. Clean-reads were mapped to *Rhinolophus sinicus* genome by hisat2. Read counts of each gene were calculated by stringtie and prepDE.py scripts. The count matrix was then processed by DESeq2 for normalization and expression profiles.

### Gene Ontology (GO) analysis

Differential genes were obtained by comparing the each macrophage cluster with others, than the genes were used for analysis with p-adjust value<0.05, mean expression value >1. The differential expression more than two times were further performed Gene ontology (GO) analysis using clusterprofiler package^58^ with mouse database (org.Mm.eg.db). A P value ≤0.05 was considered significant and enriched GO terms were sorted by counts. A column chart was plotted using top 10 GO terms.

### Comparative analysis of gene profiles in bulk and single-cell sequencing

The characteristics genes in each tissue were selected from bulk-seq data if their expression level were more than 50, and 10-fold higher than other tissues. Heat-map was made by Seaborn package (0.9.0), showing the genes expression level in selected tissue compared to the average of level in all tissues. These genes were than analyzed in single-cell sequencing data to decide the distribution in each tissue by average expression.

To compare the human and mouse gene expression with the bat respectively, we use the public databases of human metadata available on GEO (accession GSE130148, GSE134355) and mouse metadata online (http://tabula-muris.ds.czbiohub.org/) for later normalization and gene expression comparison.

### Correlations of cell type specific transcription genes

After the decision of cell types with significant transcription genes (average difference of > 1), the average gene transcription factor sets that distinguish each individual cell type from all other cells was calculated. The pearson’s correlations of specific cell types were analyzed with corr imbed in pandas package (version 0.23.4).

### T cells analysis

T cells analysis was performed by involved all organs. The differential analysis of each organ was performed, and the gene with log2 fold change ≥2 and pval_adj <1 was extracted. PCA is used for dimensionality reduction of T cells data. And then T cells were clustered by using leiden model, and reduced-dimensional mapping by using umap.

### Amino acid identity of viral receptor genes

To analyze the identity of viral receptor genes, all related coding sequences were downloaded from Ensembl and GenBank representing human, bats and mouse. The sequences were manual checked to avoid false annotation or different isoforms, then ClustalW Multiple alignment in BioEdit version 7.0.5.3 was used for amino acids sequence alignment between Chines horseshoe bat (*Rhinolophus sinicus*) and the other species.

### Single-molecule fluorescent *in situ* hybridization

Probe libraries were custom designed and constructed by Advanced Cell Diagnostics (ACD, Newark, CA) for bat SFTPC and CLDN5. The single molecule FISH probe libraries consisted of 20 probes with length of 50 bps. The probe libraries of SFTPC and CLDN5 were respectively coupled to HRP-C1 and HRP-C2, then stained with Opal™ fluorescent reagents. The single cells were washed with DPBS, fixed in 10% neutral formalin buffer for 1h at 37 □, then centrifuged at 250×g for 10 min and resuspended in 70% ethanol for incubating at RT for 10 min and stored at 4°C. Adjust the cell density to 1×10^6^ cells/ml. Cell suspension droplets were added onto the slices and dried, then incubated in 50% ethanol, 70% ethanol and 100% ethanol, for 5min at each step. The slices were dried at 37°C for 30 min, then draw for 2-4 times around the cell spot by using the hydrophobic barrier pen. The probes hybridizing was performed in accordance with the manufacturer’s instructions by using RNAscope® Multiplex Fluorescent Reagent Kit v2 and hybridization oven (HybEZ™, ACD, Newark, CA). The cells were incubated with the Hybridize Probes, hybridize Amp 1, Amp 2 and Amp 3 at 40□ for 2h, 30min, and 15min, respectively, then incubated with horseradish peroxidase (HRP)-C1 and HRP-C2. The nucleus was stained with DAPI (Invitrogen, Waltham, MA, USA) for 30 second and ProLong™ Gold antifade reagent was placed on the slices. Images were taken by using Vectra Polaris Automated Quantitative Pathology Imaging System (PerkinElmer, Waltham, MA, USA).

### Cell culture

Primary bat pulmonary fibroblast (BPF) cells were cultured from one lung of the Chinese horseshoe bat (*Rhinolophus sinicus*). The lung was pretreated followed the same procedure of single cell preparation. The cells were suspended in Roswell Park Memorial Institute (RPMI) 1640 medium (Thermo Fisher Scientific, CA, USA) containing 10% fetal bovine serum (FBS) (Hyclone, Logan, UT, USA) and 1% penicillin (10,000 IU)-streptomycin (10,000μg/mL) (PS) (Thermo Fisher Scientific, CA, USA), then cultured in a 24-well culture plate for 48 h until the fibroblasts attached to the bottom of the plate. Then the culture medium was replaced by Fibroblast Medium (ScienCell, Carlsbad, CA, USA) containing 2% FBS and 1% PS. The BPF cells were tested by mycoplasm detection kit (Lonza, Walkersville, MD, USA). The cell type was confirmed by in situ hybridization using RNAscope® Probes (ACD) targeted to fibronectin 1 (*FN1*) and asporin (*ASPN*) genes.

### Cell lines and viruses

Human Pulmonary Fibroblasts (HPFs, ScienCell, Carlsbad, CA, USA) are characterized by immunofluorescence with antibody specific to fibronectin (Santa Cruz, CA, USA) and Alexa Fluor 488-ligated second antibody (ZSGB-BIO, China). HPFs were cultured in Fibroblast Medium (ScienCell). Vesicular Stomatitis Virus (VSV) was stored in our lab and the viral titer was 5.25×10^9^ plaque forming unit (PFU) /ml.

### Quantitative reverse transcription PCR (qRT-PCR)

BPFs and HPFs were cultured in 12-well plate (2.5×10^5^ cells/well) and transfected with 1μg poly (I:C) (InvivoGen, CA, USA) or 2 μg ISD (InvivoGen, CA, USA) by lipofectamine 2000 reagent (Thermo Fisher Scientific, CA, USA), the culture medium was replaced with Opti-MEM™ Reduced Serum Medium (Thermo Fisher Scientific, NY, USA) after 4h transfection, and the cells were collected at 4 h, 8h, 12h and 24h post transfection. BPFs and HPFs were also stimulated with 10 μg/ml of poly (I:C) or 2μg/ml of R848 (MCE, NJ, USA), or medium as control. Poly (I:C) low molecular weight (LMW) and high molecular weight (HMW) (InvivoGen, CA, USA) were used initially and they would stimulate the signaling pathway in the two cells. Then the poly (I:C) HMW was then used furtherly. For VSV infection, both cells were infected with VSV at MOI of 0.5. The cells in each well were collected at 4 h, 8h, 12h and 24h post stimulation or infection. RNA was isolated as described above. Total 500 ng RNA was used to synthesize cDNA by using Moloney-murine leukemia virus (M-MLV) reverse transcriptase (Promega, Madison, WI). Diluted cDNA was used in each quantitative reverse transcription-PCR (qRT-PCR). Primers used in qRT-PCR were listed in Supplementary material. The qRT-PCR was performed by using Bio-rad with real-time CFX96 amplifier (Bio-Rad Laboratories, Inc., USA) using the TB Greeen™ Premix Ex Taq™ (TaKaRa, Japan). The primers targeted to human and bat IFNβ, IL-6, TNFα, RIG-I and MDA5 were used. Fold change expression of genes were calculated by ^ΔΔ^CT method. The mean value was from three replicates, and error bars represent standard deviations.

## Data availability

All gene expression data from single cell and bulk sequencing have deposited in the Genome Sequence Archive (GPB 2017) in National Genomics Data Center (NAR 2020), under project that is publicly accessible at https://bigd.big.ac.cn/gsa.

## Acknowledgments

We thank Dr. Zhuo Zhou (Peking University, Beijing, China) give us valuable suggestion to improve the paper. This study was funded in part by the Chinese Academy of Medical Sciences (CAMS) Innovation Fund for Medical Sciences (2016-I2M-1-014, 2019-I2M-2-001), the National Major Science & Technology Project for Control and Prevention of Major Infectious Diseases in China (2017ZX10103004), the Non-profit Central Research Institute Fund of CAMS (2019PT310029), the Ministry of Science and Technology of China (2018YFA0108100), the National Natural Science Foundation of China (21525521, 21927802), Beijing Advanced Innovation Center for Genomics, and 2018 Beijing Brain Initiative (Z181100001518004).

## Author contributions

JWW, JBW, YYH, and LLR conceived and designed experiments. CHW, YX, LLH, ZQW and XX performed the experiments. CW, JCY, LLR, AOP, LG, JBW, YYH, HPW, LLH, HZ, CHW, YWL, JCZ, LSS, MKL and XBL analyzed the data. LLR, LG, CHW, YYH, JBW, SRQ and JWW wrote the manuscript. All authors reviewed the manuscript.

## Extended Data

**Extended Data Figure 1.**
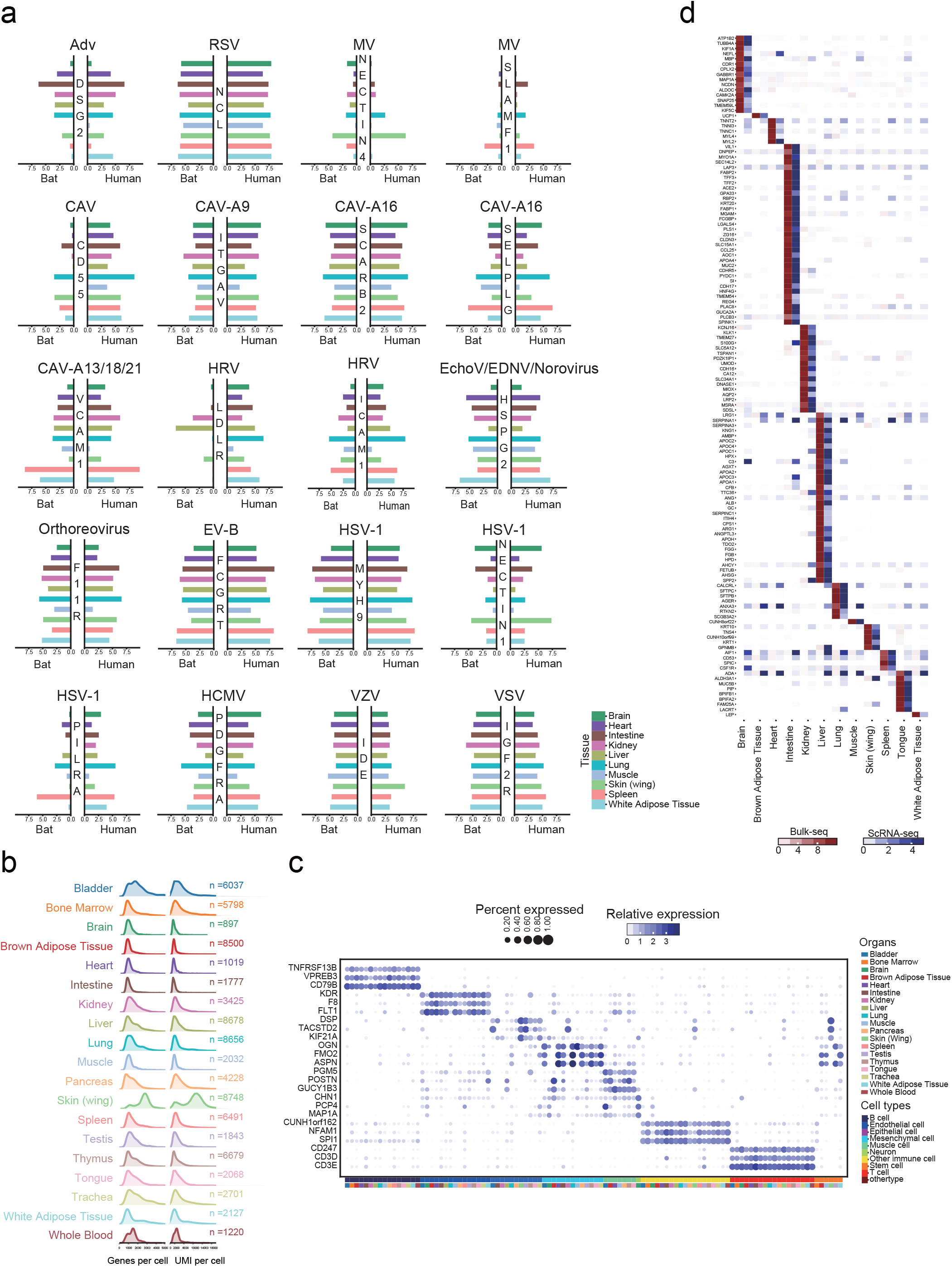
Transcriptomic analysis by bulk sequencing and the comparisons to single cell sequencing. a, Expression level of selected viral receptor genes in organs based on bulk-seq data of bat and human. Adv: Adenovirus; RSV: Respiratory syncytial virus; MV: measles virus; CAV: Coxsackie virus; CAV-A9: Coxsackie virus A9; CAV-A16: Coxsackie virus A16; CAV-A13/18/21: Coxsackie virus A13/18/21; HRV: Rhinovirus; EchoV: Echovirus; DENV: Dengue virus; EV-B: Enterovirus B; HSV-1:Herpes simplex virus; HCMV: human cytomegalovirus; VZV: Varicella zoster virus; VSV: Vesicular stomatitis virus. b, Histogram of the number of detected genes (left) and UMIs (right) per cell for each organ. c, Dot plot visualization of differentially expressed genes across clustered cells. d, The comparison of differential transcriptional genes between bulk-seq and single cell (sc)-seq data in each organ.

**Extended Data Figure 2.**
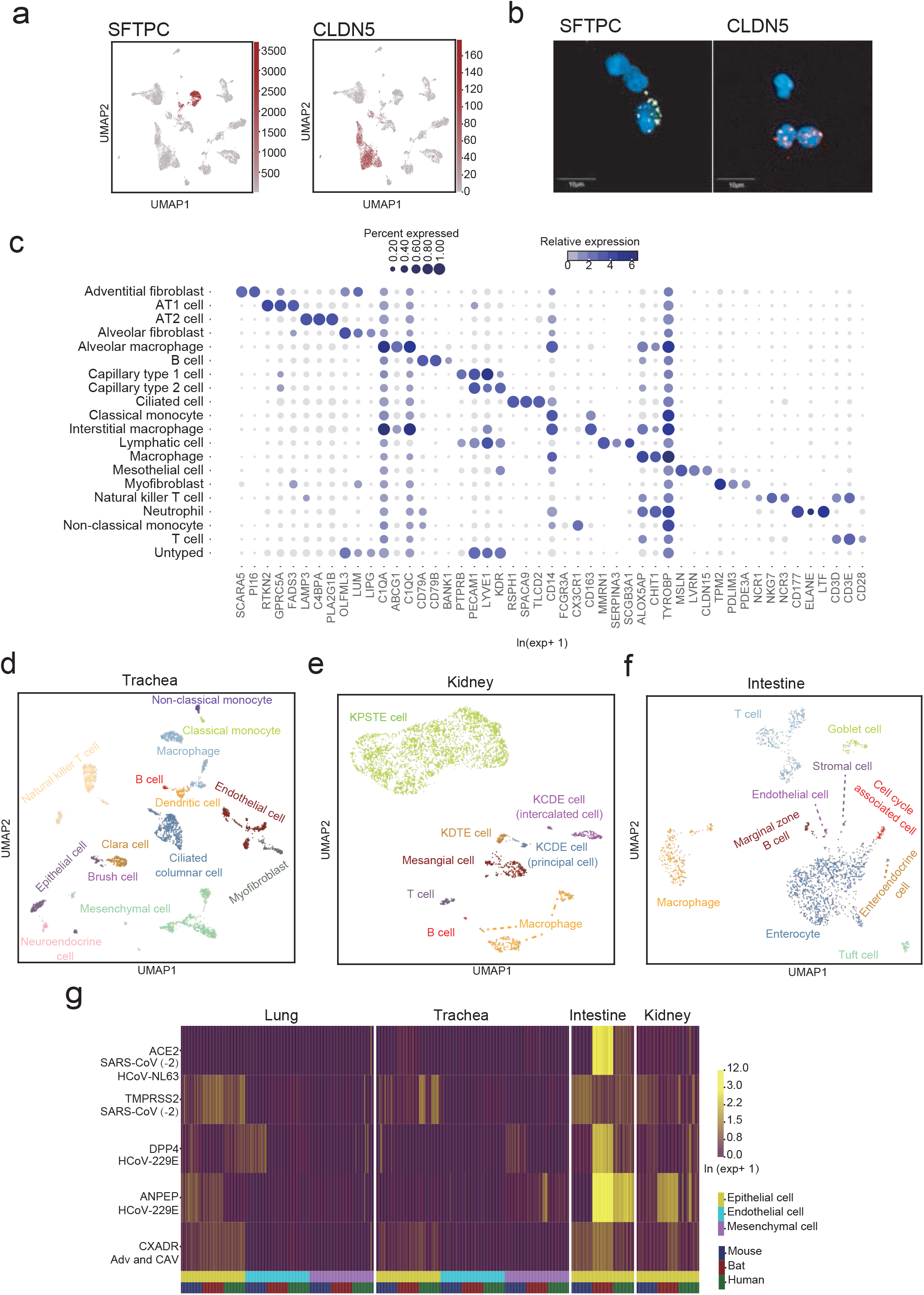
Analysis of the cell types and receptor genes distribution patterns in single cell level. a, UMAP plots of expression for genes specifically expressed in particular cell types (SFTPC in AT2 and CLDN5 in endothelial cells). Gene expression levels are indicated by shades of red. b, Single-molecule fluorescence in situ hybridization of SFTPC (Opal 520) and CLDN5 (Opal 690) on lung single cells droplet slices. c, Dot plot visualization of selected marker genes for each cell type. The size of the dot encodes the percentage of cells within a cell type in which that marker was detected, and the color encodes the average expression level. d, UMAP visualization and marker-based annotation of trachea cells. Cells are colored by cell-type. e, UMAP visualization and marker-based annotation of kidney cells. Cells are colored by cell-type. f, UMAP visualization and marker-based annotation of intestine cells. Cells are colored by cell-type. g, Comparisons of the expression patterns of the respiratory virus receptor genes, *ACE2*, *DPP4*, *ANPEP*, *CXADR* and the *TMPRSS2* in endothelial cells, epithelia cells, and mesenchymal cells in single cell levels in bat compared to that of human and mouse. Only epithelial cells in intestine and kidney were selected for the comparisons according to the available data. SARS-CoV, Severe acute respiratory syncytial virus; HCoV, Human coronavirus; Adv: Adenovirus; CAV: Coxsackie virus.

**Extended Data Figure 3.**
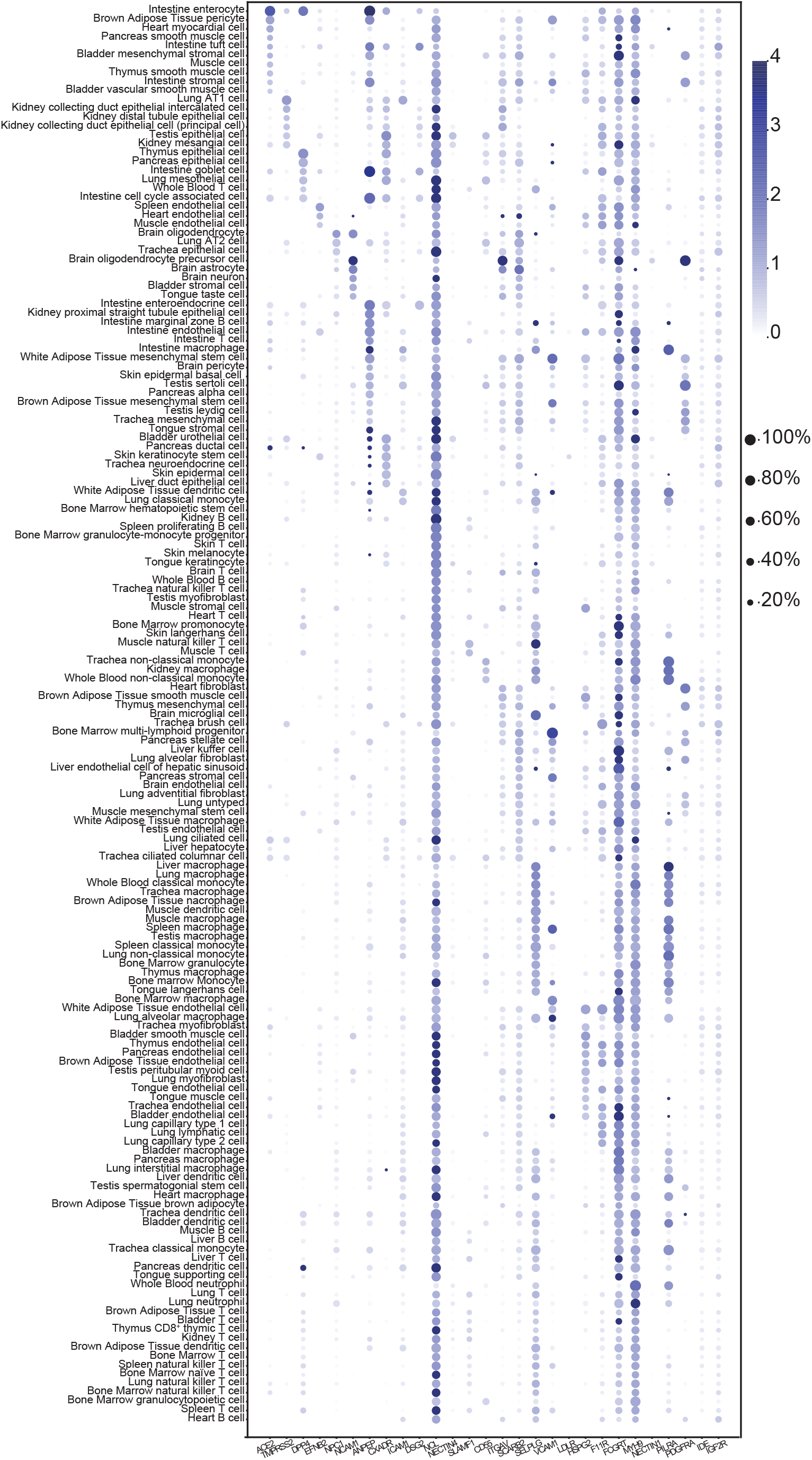
The known human viral receptors genes expressed in bat across the cell types. Dot plots of expression for viral receptor genes in featured cell types. The size of the dot encodes the percentage of cells within a cell type in which that marker was detected, and the color encodes the average expression level.

**Extended Data Figure 4.**
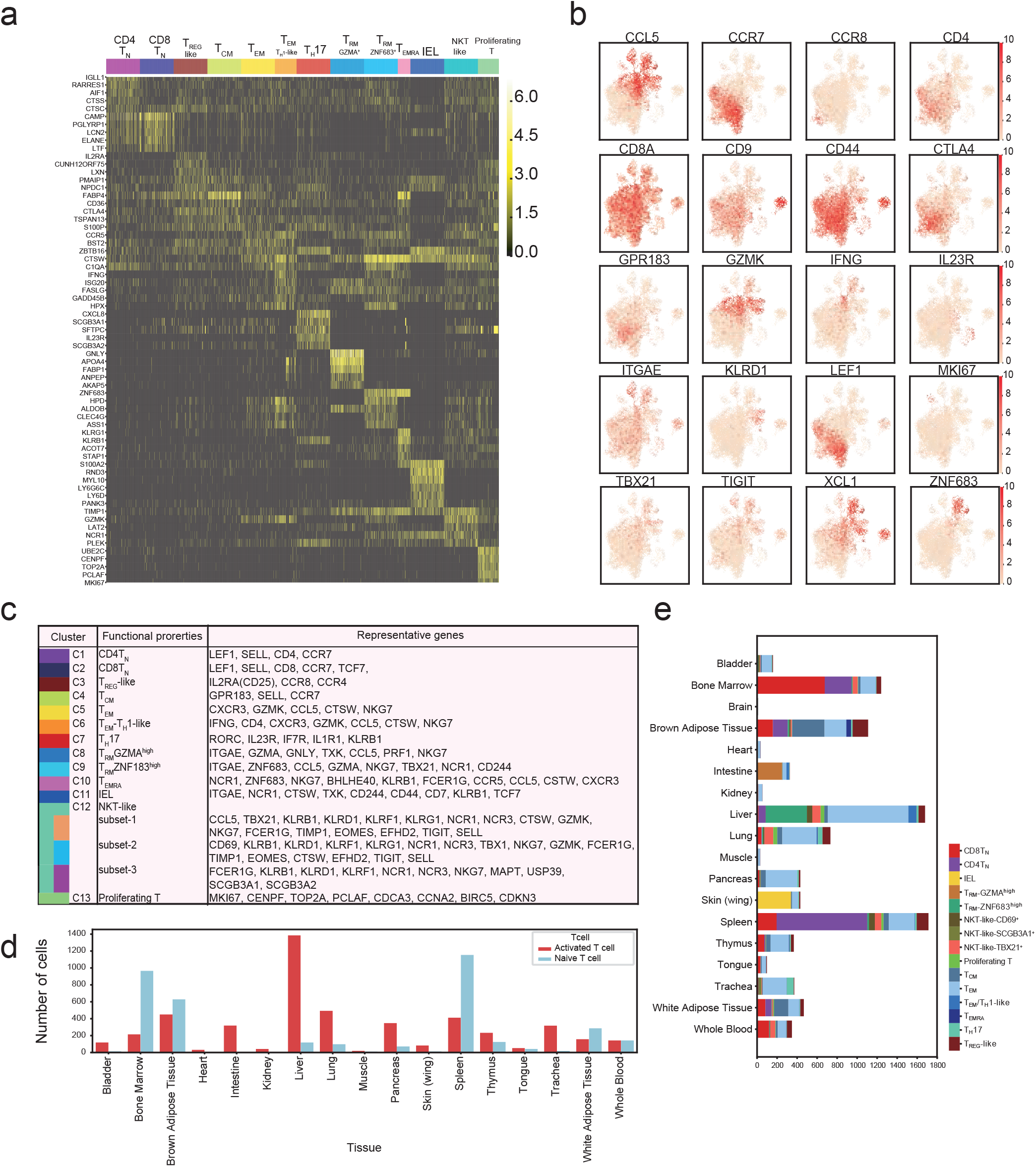
Summary of functional properties of various T cell clusters. a, Heatmap of unique signature genes for thirteen T cell clusters. Selective specifically expressed genes are marked alongside. b, UMAP plots of expression levels of selected genes in different clusters indicated by the colored oval corresponding to Figure 3a. c, Overview of T cell cluster characteristics. d, The number of activated T cells and naïve T cells in different tissue.

**Extended Data Figure 5.**
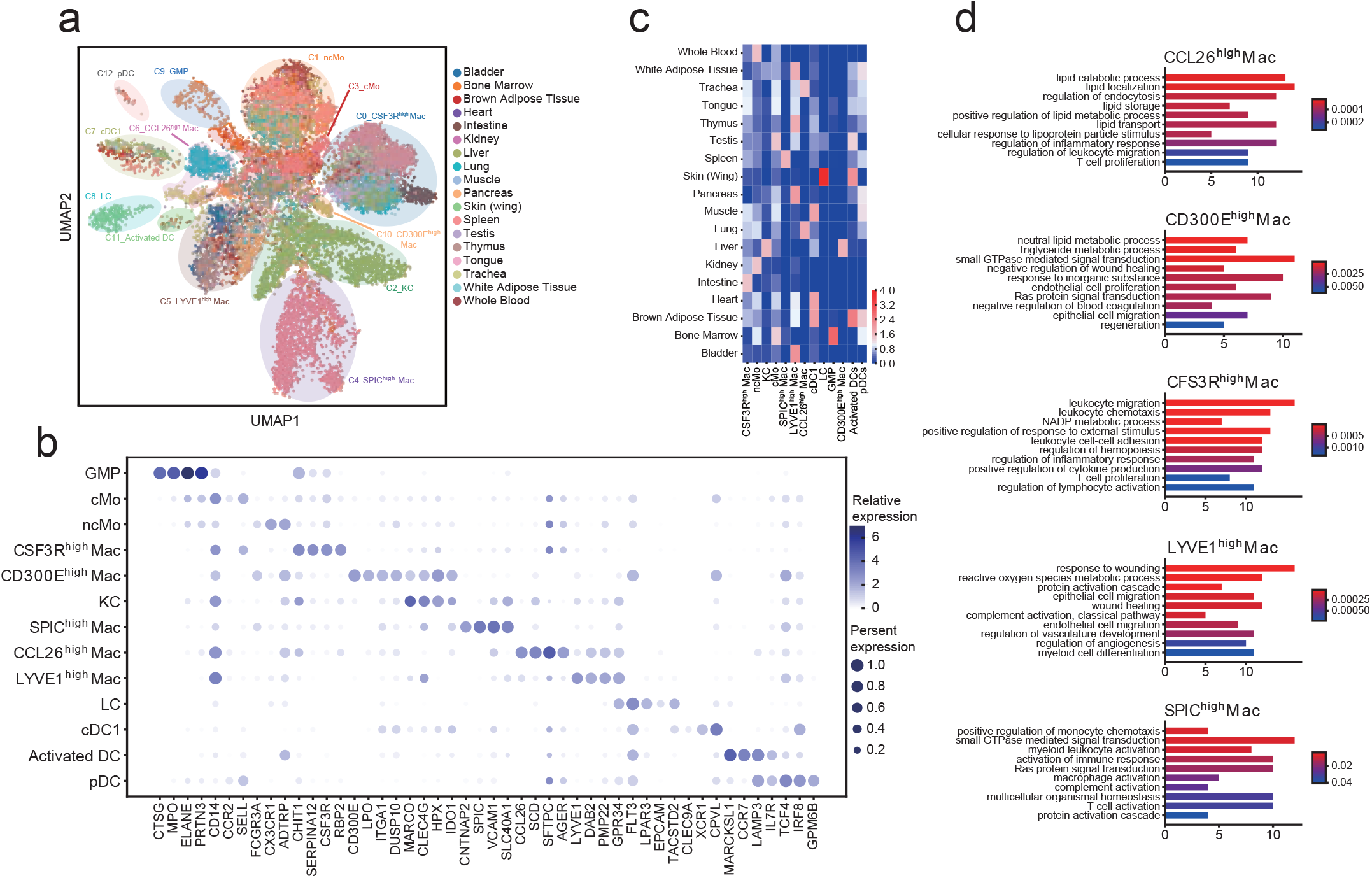
Analysis of mononuclear phagocytes. a, UMAP visualization of mononuclear phagocytes from Chinese horseshoe bats (*Rhinolophus sinicus*), showing the formation of 13 main clusters shown in different organs. The functional description of each cluster is determined by the gene expression characteristics of each cluster. b, Dot plot visualization of each cell type and selected marker gene. The size of the dot encodes the percentage of cells within a cell type in which that marker was detected, and the color encodes the average expression level. c, Tissue preference of mononuclear phagocytes estimated by proportion based on 10x data.d, Gene ontology analysis of macrophage with high expressions of *CCL26, CD300, CSF1R, LYVE1 and SPIC*.

**Extended Data Figure 6.**
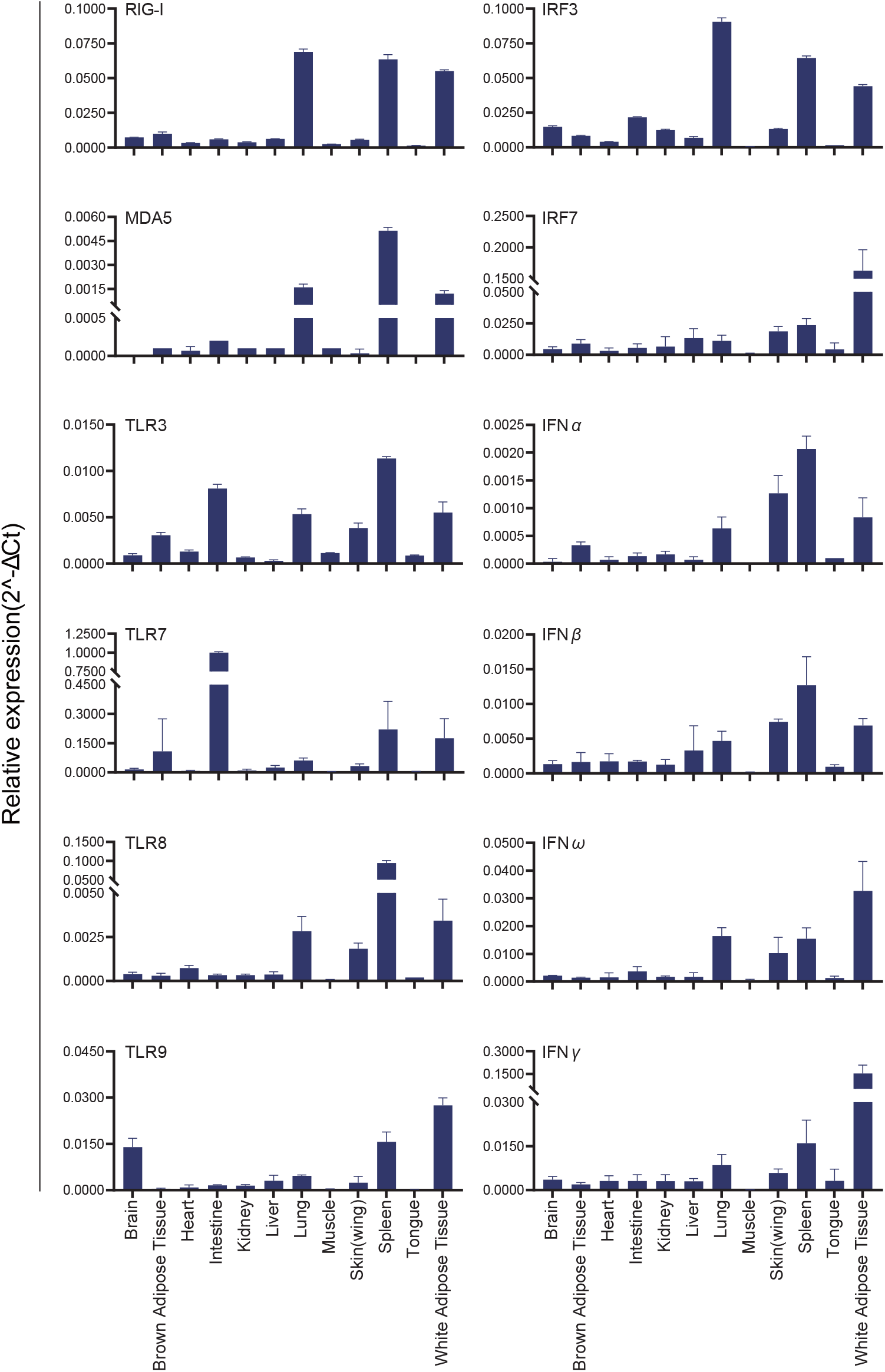
Analysis of innate immune gene mRNA expression in Chinese horseshoe bat tissues. Tissue mRNA expression levels of RIG-I, MDA5, TLR3, TLR7, TLR8, TLR9, IRF3, IRF7, IFNα, IFNβ, IFNω and IFNr were determined by qRT-PCR and normalised relative to GAPDH. Error bars represent standard deviation.

**Extended Data Figure 7.**
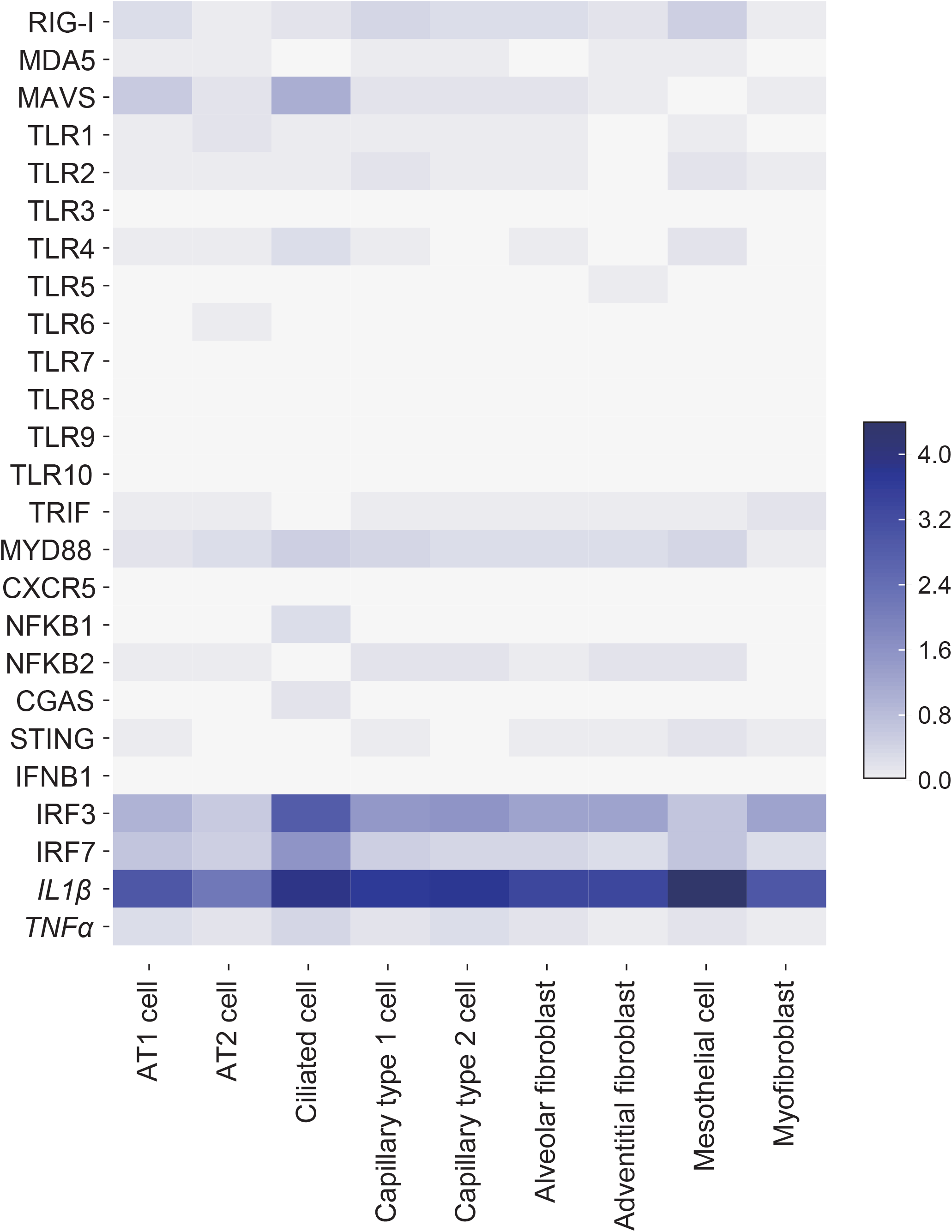
Heatmap of innate immune response genes expression in lung epithelial cells, endothelial cells and stromal cells.

## Supplementary Material

### KEY RESOURCES TABLE

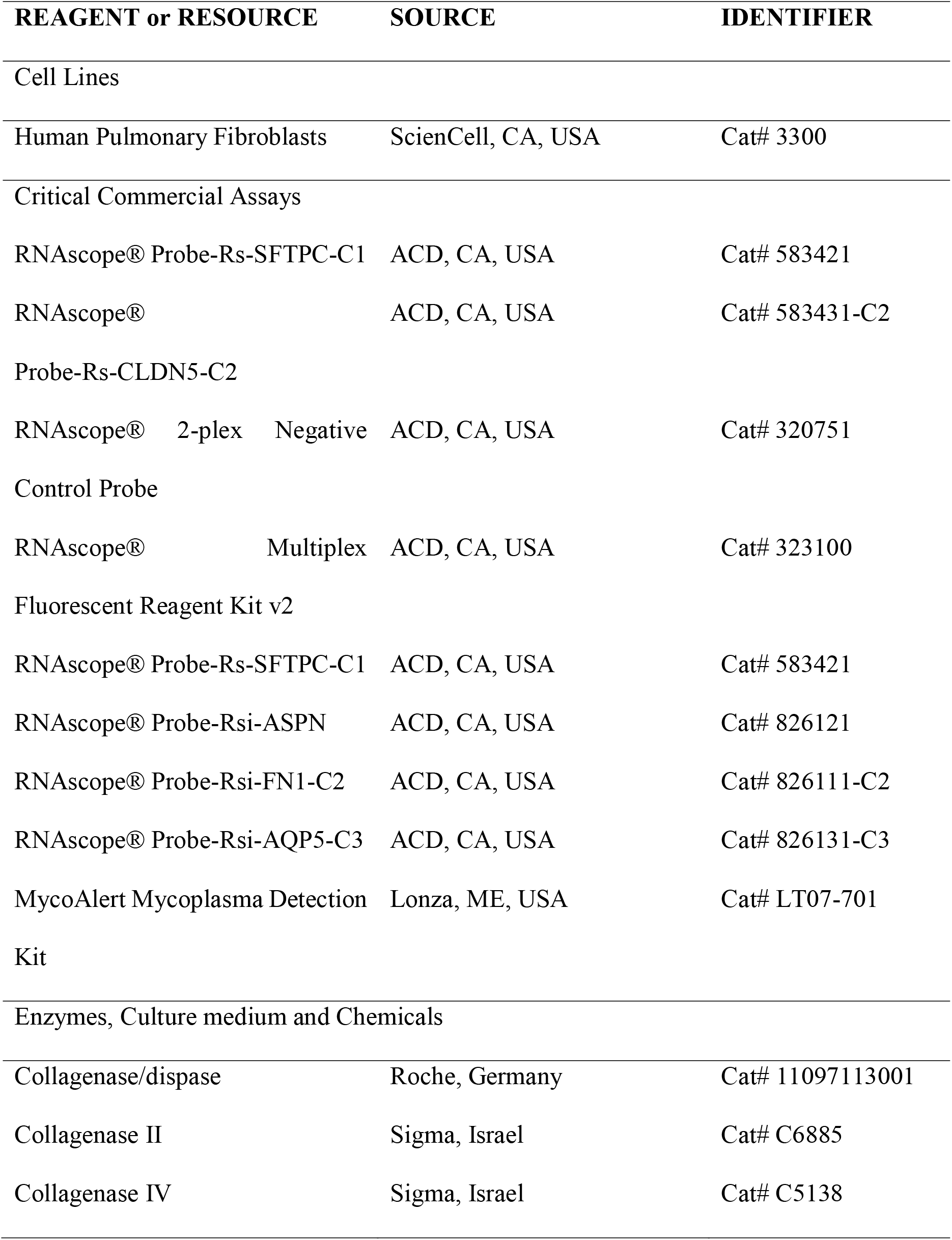

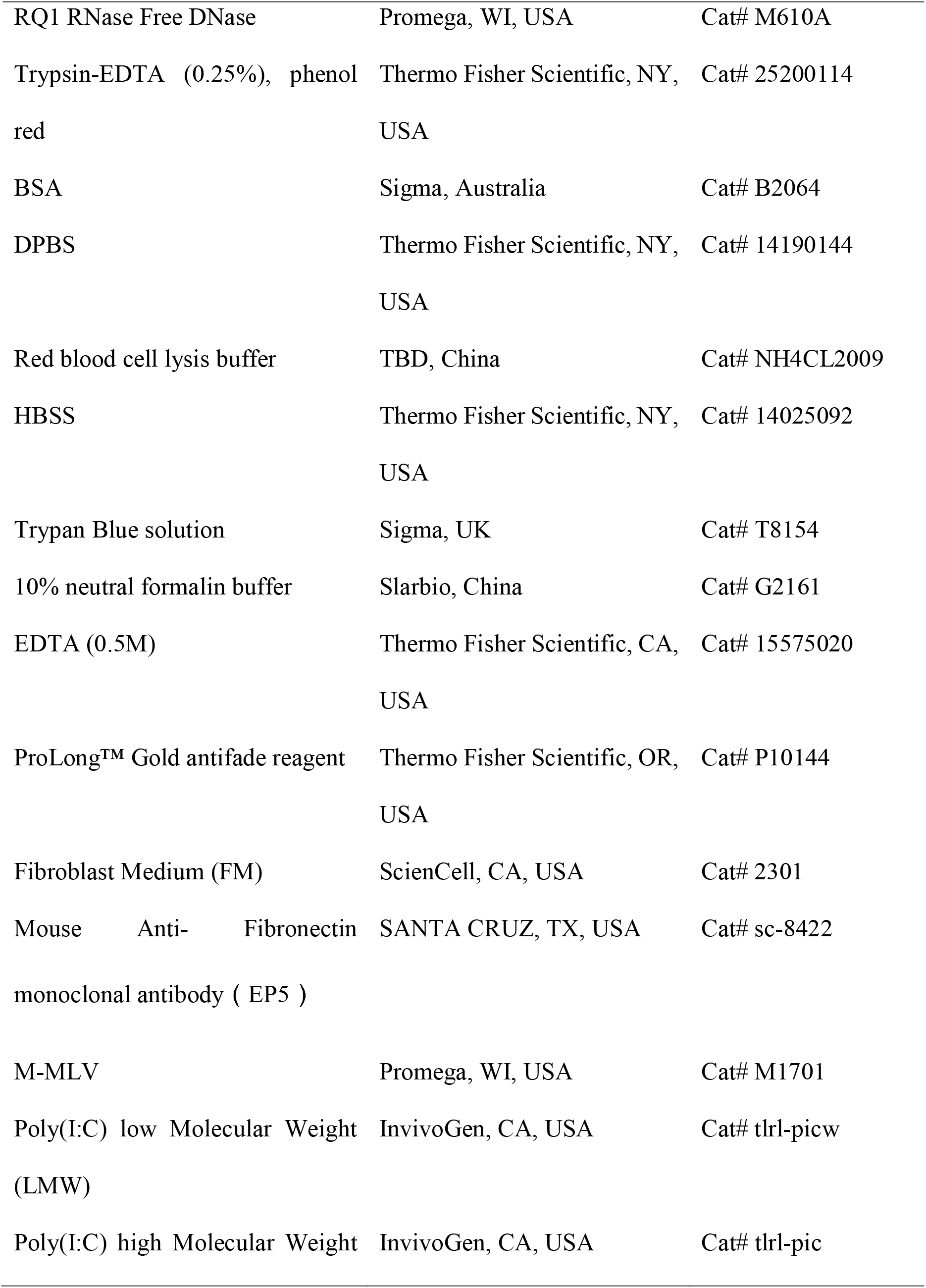

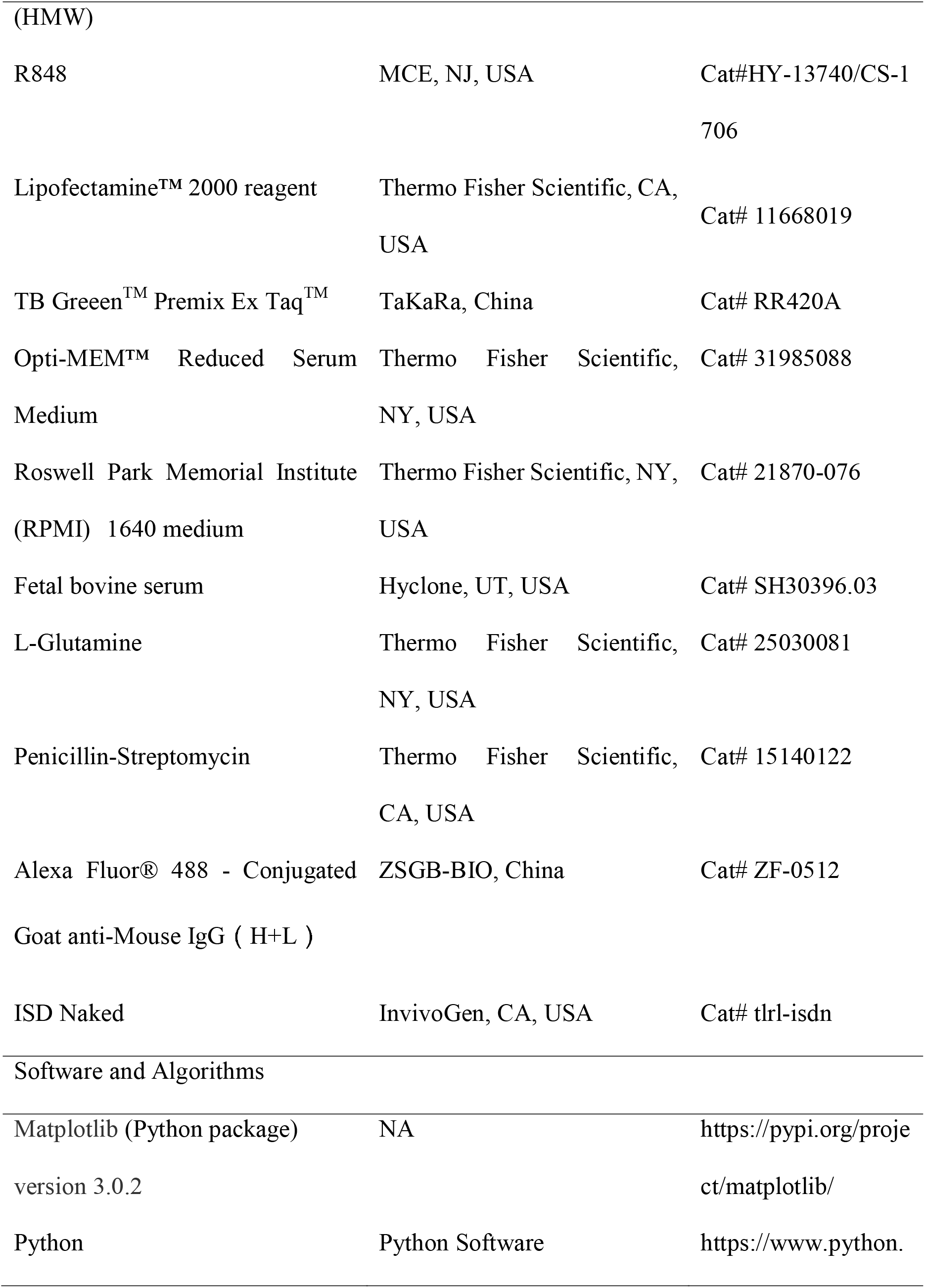

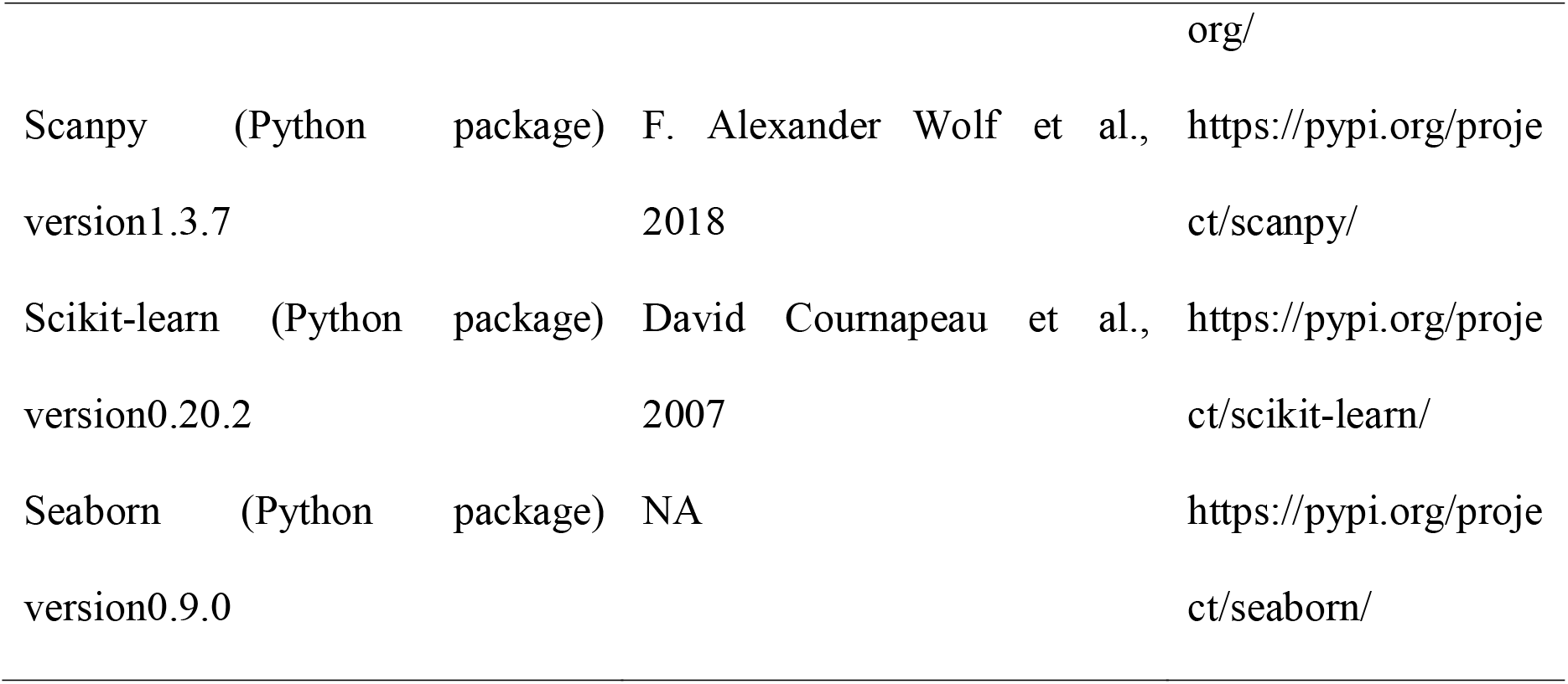

### Primers for qRT-PCR

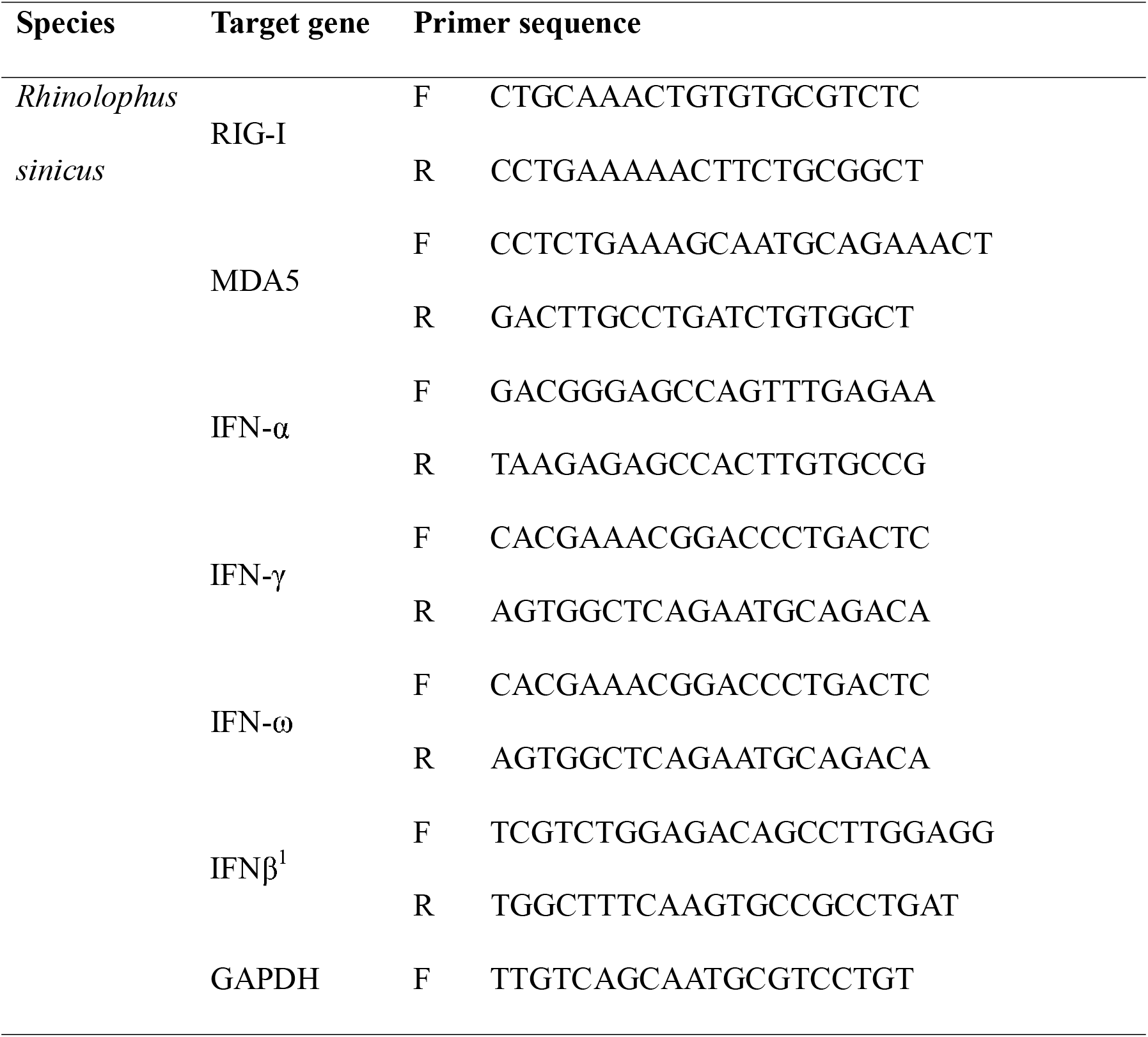

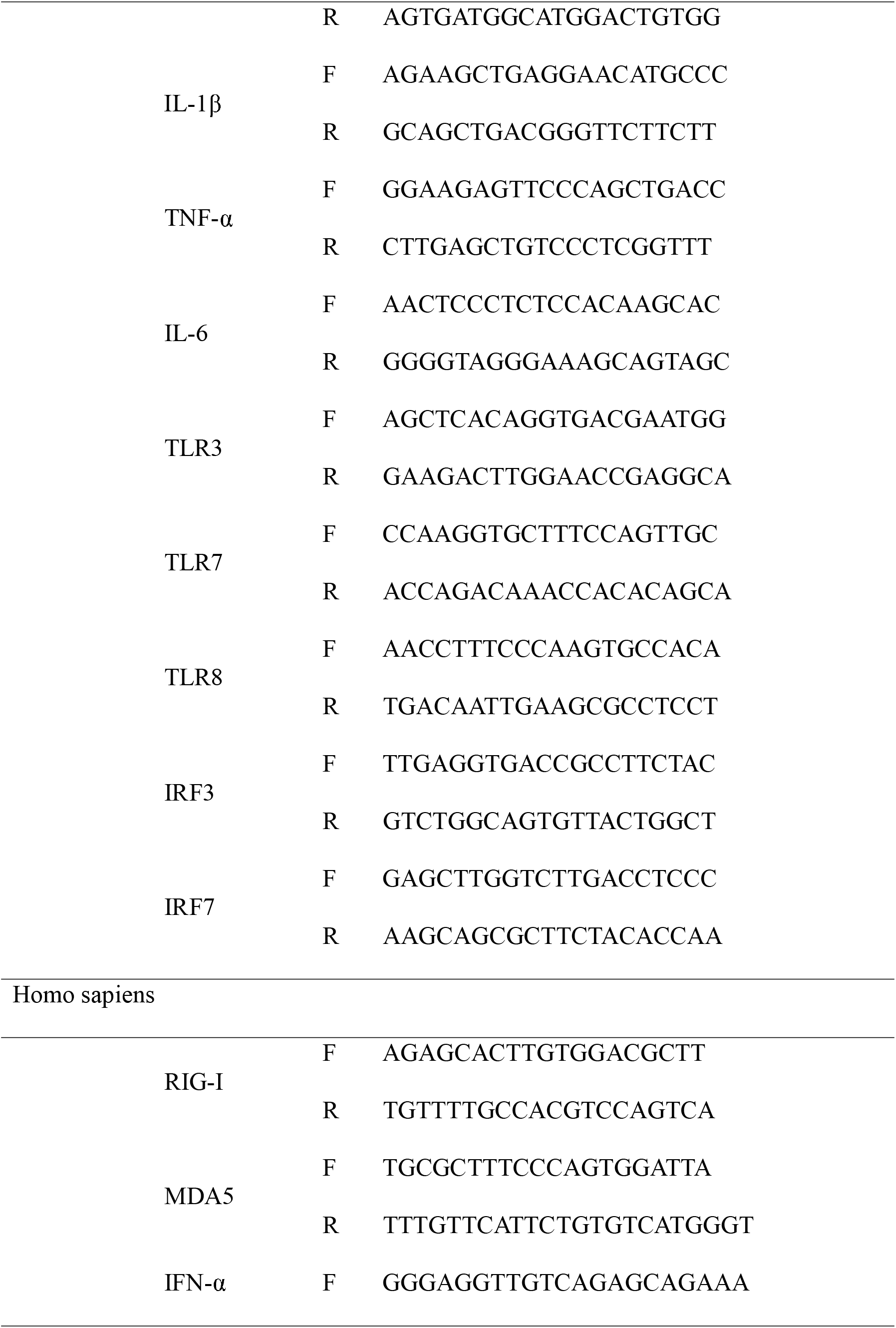

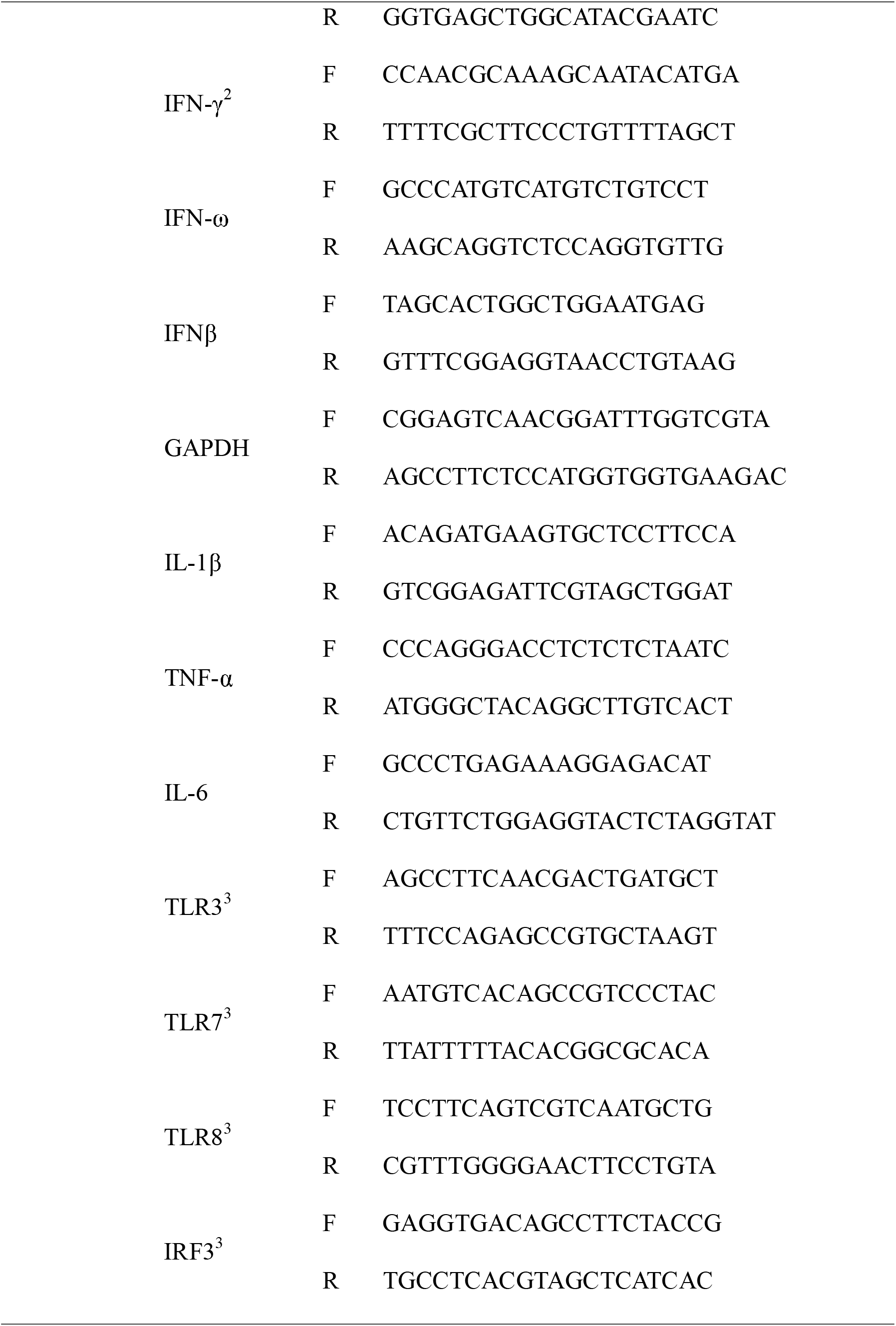

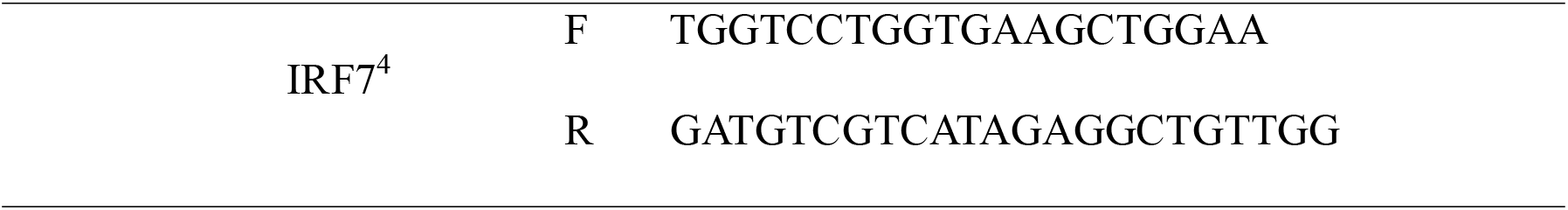

